# Silent cold-sensing neurons drive cold allodynia in neuropathic pain states

**DOI:** 10.1101/2020.05.02.073999

**Authors:** Donald Iain MacDonald, Ana P. Luiz, Queensta Millet, Edward C. Emery, John N. Wood

## Abstract

Neuropathic pain patients often experience innocuous cooling as excruciating pain. The cell and molecular basis of this cold allodynia is little understood. We used *in vivo* calcium imaging of sensory ganglia to investigate the activity of peripheral cold-sensing neurons in three mouse models of neuropathic pain: oxaliplatin-induced neuropathy, partial sciatic nerve ligation and ciguatera poisoning. In control mice, cold-sensing neurons were few in number and small in size. In neuropathic animals with cold allodynia, a set of normally silent large-diameter neurons became sensitive to cooling. Many silent cold-sensing neurons expressed the nociceptor markers Na_V_1.8 and CGRPα. Ablating these neurons diminished cold allodynia. Blocking K_V_1 voltage-gated potassium channels was sufficient to trigger *de novo* cold sensitivity in silent cold-sensing neurons. Thus silent cold-sensing neurons are unmasked in diverse neuropathic pain states and cold allodynia results from peripheral sensitization caused by altered nociceptor excitability.

**Graphical Abstract:** 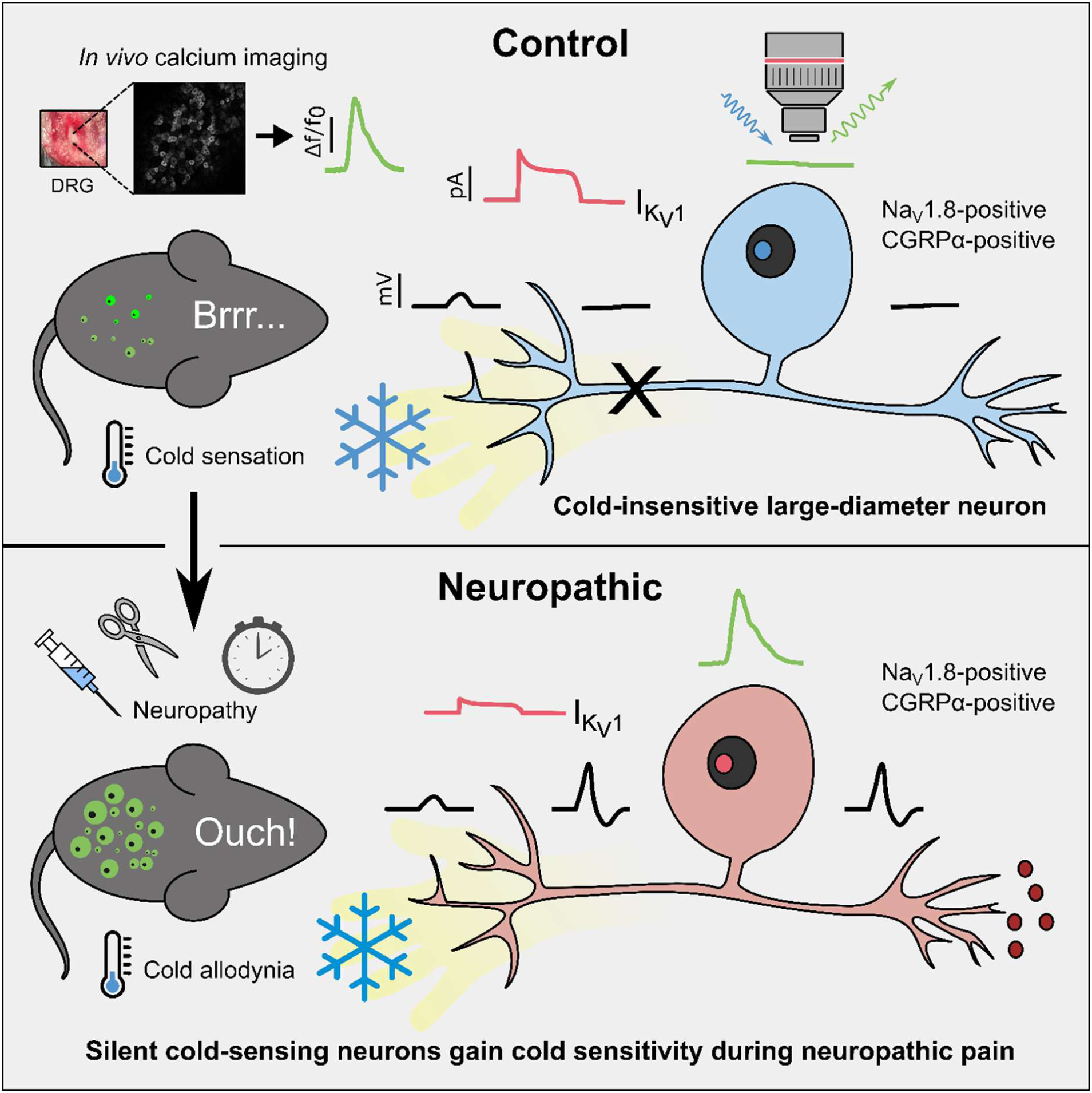

## 1 Introduction

Both pain and temperature sensation are essential for survival, but can go awry. One in five people is burdened by chronic pain and few effective treatments are available.^1^ Among the most unpleasant symptoms of chronic pain is cold allodynia, when people experience innocuous cooling as excruciating pain.^2,3^ Cold allodynia is a common complaint of patients suffering from neuropathic pain caused by chemotherapy, nerve injury or ciguatera poisoning, with a prevalence of up to 90%.^3,4^ How do neuropathic pain conditions with different etiologies give rise to the same sensory disturbances of cold-evoked pain? Despite advances in our understanding of cold sensation during the healthy state, the pathophysiological mechanisms underlying cold allodynia remain elusive.

Chronic pain has been ascribed to both peripheral and central sensitization of the sensory pathways subserving nociception and pain.^5,6^ Is cold-evoked neuropathic pain peripheral in origin? Under physiological conditions, peripheral sensory neurons show modality-specific responses to cold, with ‘labelled lines’ for both mild and extreme cooling.^7–12^ Cold detection depends on cold-gated ion channels like Trpm8, as well as the expression of different sodium and potassium channels that control excitability at low temperatures.^13–20^ Studies of ion channel knockout mice suggest cold allodynia is maintained by canonical TRP channels and potassium channels expressed by unmyelinated C fibres.^8,14,21–27^ A role for sodium channels enriched in A fibres is also evident, however.^23,28–30^ Mechanistic investigation of the cells and molecules driving cold allodynia has so far proved difficult because of the challenge in recording large numbers of cold-responsive afferents, as well as the limitations of traditional cold pain behaviour tests.^31^

To directly investigate if cold allodynia results from plasticity in peripheral sensory neurons, we used *in vivo* calcium imaging to explore how the activity of cold-sensing neurons is altered in neuropathic pain. Here we identify a previously undescribed set of large-diameter silent cold-sensing neurons that contribute to cold allodynia in diverse neuropathic pain states and provide evidence for the involvement of K_V_1 potassium channels in unmasking their latent cold sensitivity.

## 2 Results

### 2.1 Silent cold-sensing neurons are unmasked during chemotherapy-induced neuropathy

To investigate the mechanisms of cold allodynia, we used *in vivo* calcium imaging to explore how sensory neuron responses to cooling are altered in different mouse models of neuropathic pain. We focused initially on chemotherapy-induced neuropathy because ∼90% of cancer patients treated with the chemotherapeutic drug oxaliplatin develop cold allodynia.^3,4^ Pirt-GCaMP3 mice expressing GCaMP3 in all sensory neurons were treated with oxaliplatin (80 µg/40 µl by hindpaw intraplantar injection). As previously reported, three hours after injection mice displayed extreme cold hypersensitivity, as measured by the number of nociceptive behaviours (lifting, licking, guarding or shaking of the ipsilateral hindpaw) when the animal was placed on a 5 °C Cold Plate (**Figure 1A**).^29^ The short-latency cold hypersensitivity observed after a single clinical dose of oxaliplatin (∼3mg/kg) in this model mimics the rapid onset of cold allodynia in patients.^29^ Oxaliplatin-injected mice developed mechanical hypersensitivity, but not heat hyperalgesia, consistent with a neuropathic state (**Figure 1A**).

**Figure 1.**
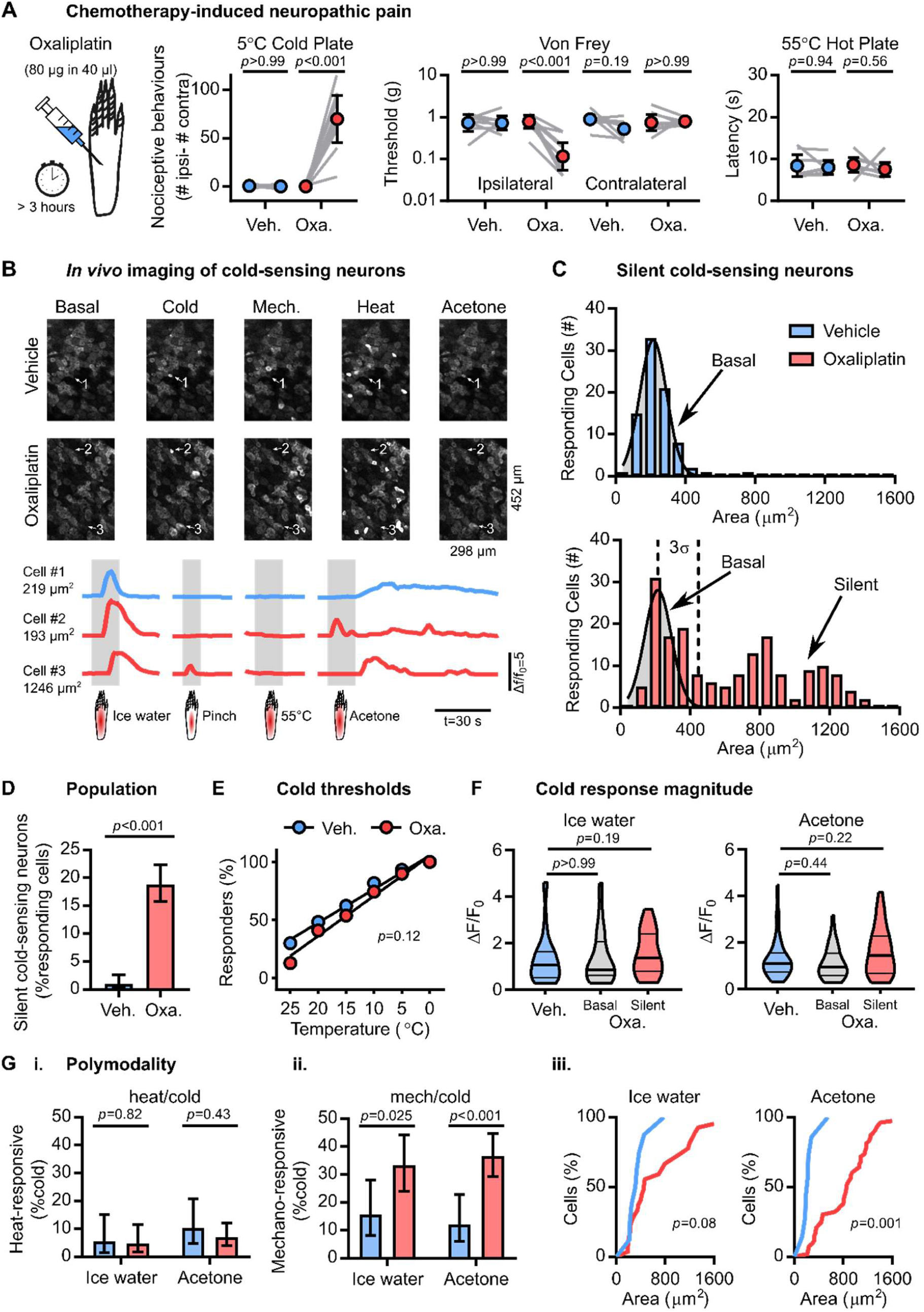
Silent cold-sensing neurons are activated by oxaliplatin-induced neuropathy. (A) Behavioural testing of the effect of oxaliplatin on different sensory modalities (cold, mechanical and heat). n=8 for vehicle and n=9 for oxaliplatin. Mean values before and after treatment were compared using repeated measures 2-way ANOVA followed by post-hoc Sidak’s test. Error bars denote 95% confidence interval. (B) Example images and traces of cold-responding neurons in vehicle- and oxaliplatin-treated animals expressing GCaMP3. Cell #1 is a small-diameter cold-sensing neuron in the vehicle condition, Cell #2 is a small-diameter basal cold-sensing neuron after oxaliplatin and Cell #3 is a large-diameter silent cold-sensing neuron unmasked by oxaliplatin that also responds to noxious mechanical stimuli (C) Histograms of cross-sectional area of all neurons responding to any cold stimulus in vehicle (top, blue, n=82) and oxaliplatin (bottom, red, n=179) groups. The distribution of areas for vehicle was fit by non-linear regression (least squares Gaussian; Bin width is 80 μm^2^; Mean=214.9 μm^2^, Std. Dev (s)=77.29 μm^2^). This model is plotted over the oxaliplatin data to aid comparison with the dashed line denoting three standard deviations from the mean. The different in the distribution of areas between groups was assessed by Kolmogorov-Smirnov test (*p*<0.001). (D) Bar plot of the percentage of responding neurons classed as silent cold-sensing neurons in vehicle and oxaliplatin groups. Proportions were compared using a χ^2^ test, and error bars denote 95% confidence intervals. (E) Relationship between the number of basal cold-sensing neurons and the drop in temperature can be fit by linear regression for both groups. For vehicle, y = −2.883 ∗ *x* + 105.2, r^2^=0.9809, n=87. For basal cold-sensing neurons afer oxaliplatin, y = −3.443 ∗ *x* + 105, r^2^=0.9802, n=39. The slopes are not significantly different (*p*=0.12). (F) Violin plots showing the peak responses evoked by cold stimuli in the vehicle group and separately in the basal and silent cold-sensing neurons from the oxaliplatin group. Ice water: n=51 for vehicle, n=40 for basal, and n=41 for silent. Acetone: n=58 for vehicle, n=57 for basal, and n=88 for silent. Medians were compared by Kruskall-Wallis test followed by Dunn’s multiple comparison’s test. (G) Quantification of the proportion of cold-sensing neurons responding to either heat (i.) or mechanical (ii.) stimuli in vehicle and oxaliplatin groups. The proportion of polymodal neurons was compared using χ^2^ test, and error bars denote confidence intervals. Ice water: n_veh_=51, n_oxa_=81. Acetone: n_veh_=58, n_oxa_=145. (iii.) Cumulative probability plots showing mechano-cold neurons have larger cross-sectional areas in the oxaliplatin group, as determined by the Kolmogorov-Smirnov test. Ice water: n_veh_=8, n_oxa_=62. Acetone: n_veh_=7, n_oxa_=53. For this experiment, 383 neurons responding to any stimulus were recorded in 8 vehicle-treated mice and 542 cells were recorded from 9 oxaliplatin-treated animals.

Using laser-scanning confocal microscopy, we imaged cold-evoked calcium signals in sensory neuron somata within the L4 dorsal root ganglion of animals treated with vehicle or oxaliplatin. There was a dramatic change in the peripheral representation of cold following oxaliplatin treatment (**Figure 1B**). In vehicle-treated mice, cold-sensing neurons were sparse and had small diameters, in agreement with earlier studies.^11,20,32^ When we measured the size of neurons responding to either ice-water or acetone applied to the glabrous skin, cross-sectional areas were normally distributed and had a mean value of 214.9 µm^2^ (**Figure 1C**). In oxaliplatin-treated animals, small cells also responded to cold, however a novel, normally cold-insensitive population of large diameter neurons also became activated by cooling. We consequently divided cold-sensing neurons from the oxaliplatin-treated group into a basal population (within three standard deviations of the mean cross-sectional area for the vehicle group), and an unmasked population (greater than three standard deviations away from this mean, >446.77 µm2) (**Figure 1C & Figure S1A**). Because these large neurons essentially never respond to cooling under physiological conditions but gained a *de novo* sensitivity to cold following oxaliplatin, we named them ‘silent cold-sensing neurons.’ The percentage of responding cells classified as silent cold-sensing neurons rose from 1% (4/383) in vehicle-to 19% (102/542) in oxaliplatin-treated animals (**Figure 1D**).

Interestingly, the response of many silent cold-sensing neurons to acetone was temporally-delayed and continued for tens of seconds beyond the initial delivery of the stimulus (**Figure 1B**). Consistent with this, when oxaliplatin-treated animals were tested with acetone, the animals showed nociceptive behavior localized to the ipsilateral paw for nearly the entire minute following acetone stimulation (**Figure S1B**).

*In vitro* experiments suggest that cold allodynia results from a shift in cellular activation thresholds such that neurons signalling extreme cold are now active at higher temperatures.^27^ In contrast, oxaliplatin did not affect thermal activation thresholds of basal cold-sensing neurons *in vivo*, when the hindpaw was stimulated with a range of temperature drops delivered by a Peltier-controlled thermode (**Figure 1E**).^20^ Moreover, when we quantified peak fluorescence intensity in response to cold as a surrogate for excitability before and after oxaliplatin, cold-evoked fluorescence intensity in both the basal and silent populations in the oxaliplatin group was no different to vehicle (**Figure 1F**). Collectively, these data indicate oxaliplatin does not markedly affect the activation thresholds or excitability of the basally-active cold-sensing neurons.

What effect did oxaliplatin have on neurons responding to other sensory modalities? There was no marked change in cross-sectional area for mechanically-sensitive neurons, but heat-activated cells showed a small shift towards larger cells (**Figure S1A**). Importantly, there was no increase in the number of cells responding to these modalities (**Figure S1C**). In addition, the magnitude of responses to mechanical stimuli was not altered, although the response to heat was reduced (**Figure S1D**). Thus oxaliplatin treatment results in a modality-specific expansion in the peripheral representation of cold through the recruitment of silent cold-sensing neurons.

What is the functional identity of silent cold-sensing neurons? We have previously shown that nociceptor polymodality is increased by treatment with inflammatory mediators that cause pain.^12^ Interestingly, there were visibly more polymodal neurons in the oxaliplatin-treated group (**Figure 1B**). Oxaliplatin increased the proportion of mechano-heat neurons from 19% (26/136) to 32% (61/193) (**Figure S1E**), which tallies with the mechanical hypersensitivity observed in these animals (**Figure 1A**). Very few cold-sensing neurons responded to heat in both oxaliplatin and vehicle conditions (**Figure 1Gi**). By contrast, oxaliplatin markedly increased the prevalence of cold-sensing neurons also responding to mechanical stimuli (**Figure 1Gii**). For ice-water, this rose from 16% (8/51) to 33% (27/81) and for acetone from 12% (7/58) to 37% (53/145). Note this number is likely an under-estimate because the application area of pinch stimuli is less than that of ice-water or acetone and thus targets a smaller receptive field. Importantly, these mechano-cold neurons were chiefly large-diameter silent cold-sensing neurons (**Figure 1Giii**). Many silent cold-sensing neurons therefore also respond to noxious mechanical stimuli, consistent with a functional identity as nociceptors.

### 2.2 Silent cold-sensing neurons are unmasked during peripheral nerve injury

Are silent cold-sensing neurons recruited in other neuropathic pain states? To mimic chronic neuropathic pain associated with nerve injury, we performed partial sciatic nerve ligation (PNL) on Pirt-GCaMP3 mice (**Figure 2A**). Two weeks after surgery, nerve injured animals developed mechanical, but not cold, hypersensitivity. At four weeks, we saw both mechanical and a modest cold hypersensitivity, but no difference in heat nociception. We therefore performed *in vivo* imaging of both nerve-injured and sham-operated mice between 4 and 5 weeks post-surgery, when cold allodynia was present (**Figure 2B**). As expected, in sham-operated mice, cold-sensing neurons were small in size with a mean area of 222.7 µm^2^ (**Figure 2C**). But after nerve injury, a set of normally cold-insensitive, large-diameter neurons began to respond to cooling (**Figure 2C**). Neurons with cross-sectional areas greater than three standard deviations away from the mean sham value (>405.4 µm^2^) were again classified as silent cold-sensing neurons. The recruitment of large cells by cold after nerve injury was apparent for both ice-water and acetone stimuli (**Figure S2A**). Consistent with the less profound behavioural cold hypersensitivity, silent cold-sensing neurons were fewer in number in nerve injured animals compared to the effect of oxaliplatin treatment. The silent cold-sensing neuron population expanded from 2% (7/373) to 15% (45/291). (**Figure 2D**).

**Figure 2.**
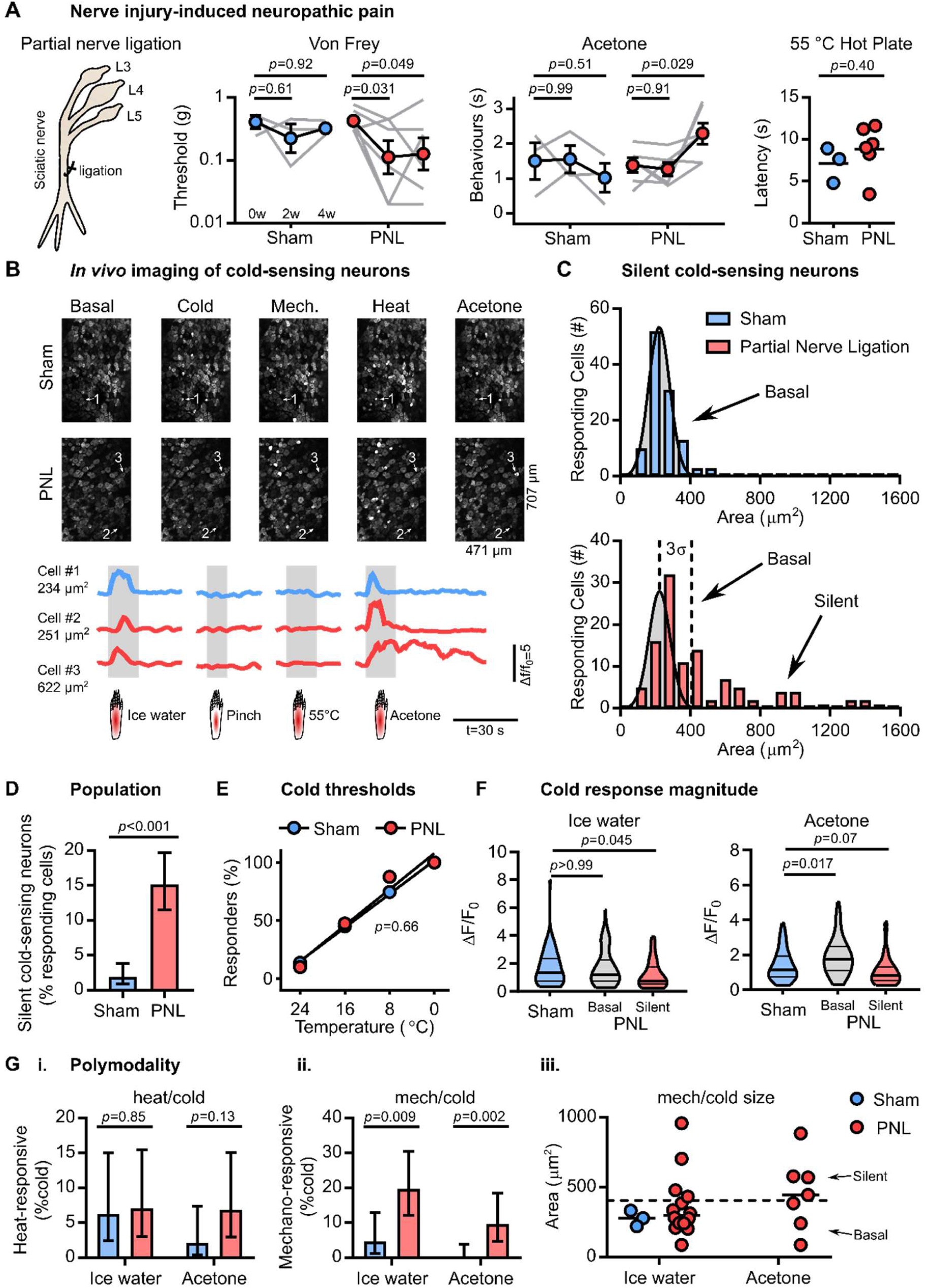
Silent cold sensing neurons are activated after partial sciatic nerve ligation. (A) Behavioural testing of the effect of partial sciatic nerve ligation (PNL) on different sensory modalities. n=3 for sham and n=6 for PNL. For Von Frey and Acetone test, means over time were compared using repeated measures 2-way ANOVA followed by post-hoc Sidak’s test. Hot Plate latencies at 4 weeks were compared using unpaired t test. Error bars denote 95% confidence interval. (B) Example images and traces of cold-responding neurons in sham- and PNL-operated animals expressing GCaMP3. Cell #1 is a small-diameter cold-sensing neuron in the sham condition, Cell #2 is a small-diameter basal cold-sensing neuron after PNL and Cell #3 is a large-diameter silent cold-sensing neuron unmasked by PNL. (C) Histograms of cross-sectional area of all neurons responding to any cold stimulus in sham (top, blue, n=113) and PNL (bottom, red, n=109) groups. The distribution of areas for sham was fit by non-linear regression (least squares Gaussian; Bin width is 80 µm^2^; Mean=222.7 µm^2^, Std. Dev. 60.9 µm^2^). This model is plotted over the PNL data to aid comparison with the dashed line denoting three standard deviations from the mean. The different in the distribution of areas between groups was assessed by Kolmogorov-Smirnov test (*p*<0.001). (D) Bar plot of the percentage of responding neurons classed as silent cold-sensing neurons in sham and PNL groups. Proportions were compared using a χ^2^ test, and error bars denote confidence intervals. (E) Relationship between the number of basal cold-sensing neurons and the drop in temperature can be fit by linear regression for both groups. For sham, y = −3.603 ∗ *x* + 101.6, r^2^=0.9979, n=51. For PNL, y = −3.875 ∗ *x* + 107.8, r^2^=0.9598, n=40. The slopes are not significantly different (*p*=0.66). (F) Violin plots showing the peak responses evoked by cold stimuli in the sham group and separately in the basal and silent cold-sensing neurons for the PNL group. Ice water: n=64 for vehicle, n=46 for basal, and n=25 for silent. Acetone: n=95 for sham, n=42 for basal, and n=31 for silent. Medians were compared by Kruskall-Wallis test followed by Dunn’s multiple comparison’s test. (G) Quantification of the proportion of cold-sensing neurons responding to either heat (i.) or mechanical (ii.) stimuli in sham and PNL groups. The proportion of polymodal neurons was compared using χ^2^ test, and error bars denote confidence intervals. Ice-water: n_sham_=64, n_PNL_=71. Acetone: n_sham_=95, n_PNL_=73. (iii.) Scatter plots showing mechano-cold neurons have both small and large cross-sectional areas in the PNL group. Mech./ice water: n_sham_=3, n_PNL_=14. Mech./acetone: n_sham_=0, n_PNL_=7. For this experiment, 373 neurons responding to any stimulus were recorded in 3 sham-operated mice and 291 cells were recorded from 6 PNL-operated animals.

As with oxaliplatin, there was no detectable shift in the thermal activation thresholds of basal cold-sensing neurons after nerve injury (**Figure 2E**). The impact of nerve injury on the excitability of cold-sensing neurons was complex. Acetone-evoked activity was enhanced in the basal population, while silent cells showed reduced responses to ice-water stimuli (**Figure 2F**). There was no change in the prevalence of heat/cold polymodal neurons (**Figure 2Gi**), however the proportion of mechano-cold cells was significantly increased (**Figure 2Gii**). For ice-water, this rose from 5% (3/64) to 20% (14/71) and for acetone from 0% (0/95) to 10% (7/73). Mechano-cold cells comprised both the basal and silent cold-sensing neurons, based on cell size (**Figure 2Giii**). Thus, the effects of nerve injury on the peripheral representation of cold is qualitatively similar to the effect of oxaliplatin, characterized by an unmasking of silent cold-sensing neurons that also often respond to noxious mechanical stimuli.

Nerve injury had variable effects on the neurons responding to other modalities. There was no change in the cell area distribution for mechanical stimuli, but the heat-activated population was larger in size (**Figure S2A**). By contrast, significantly more neurons responded to pinch (38% vs. 28%), mirroring behavioural hypersensitivity to mechanical stimuli, and there was a trend towards fewer responses to heat (40% vs. 47%) (**Figure S2B**). We saw no difference in the intensity of the response to noxious heat, but pinch-evoked peak activity was decreased (**Figure S2C**). Unlike oxaliplatin, there was no enhancement of mechano-heat polymodality (**Figure S2D**). Taken together, although nerve injury had different effects on other modalities versus oxaliplatin, the consequences for peripheral cold sensation were comparable.

### 2.3 Silent cold-sensing neurons are unmasked during ciguatera poisoning

Both the oxaliplatin and nerve injury model show a delayed onset of cold hypersensitivity. As our imaging preparation is terminal, we could not follow mice in real-time to determine if silent cold-sensing neurons are truly silent at the naive state. To induce cold allodynia within the same imaging session, we therefore turned to a mouse model of ciguatera poisoning, a marine toxin-induced neuropathy characterized by cold pain in the extremities that results from consuming contaminated seafood.^23^ Intraplantar injection of ciguatoxin-2 (P-CTX-2, 100 nM) into the hindpaw evoked cold pain by 30 minutes, as judged by both the acetone test and the unilateral cold plate at 10°C (**Figure 3A**).

**Figure 3.**
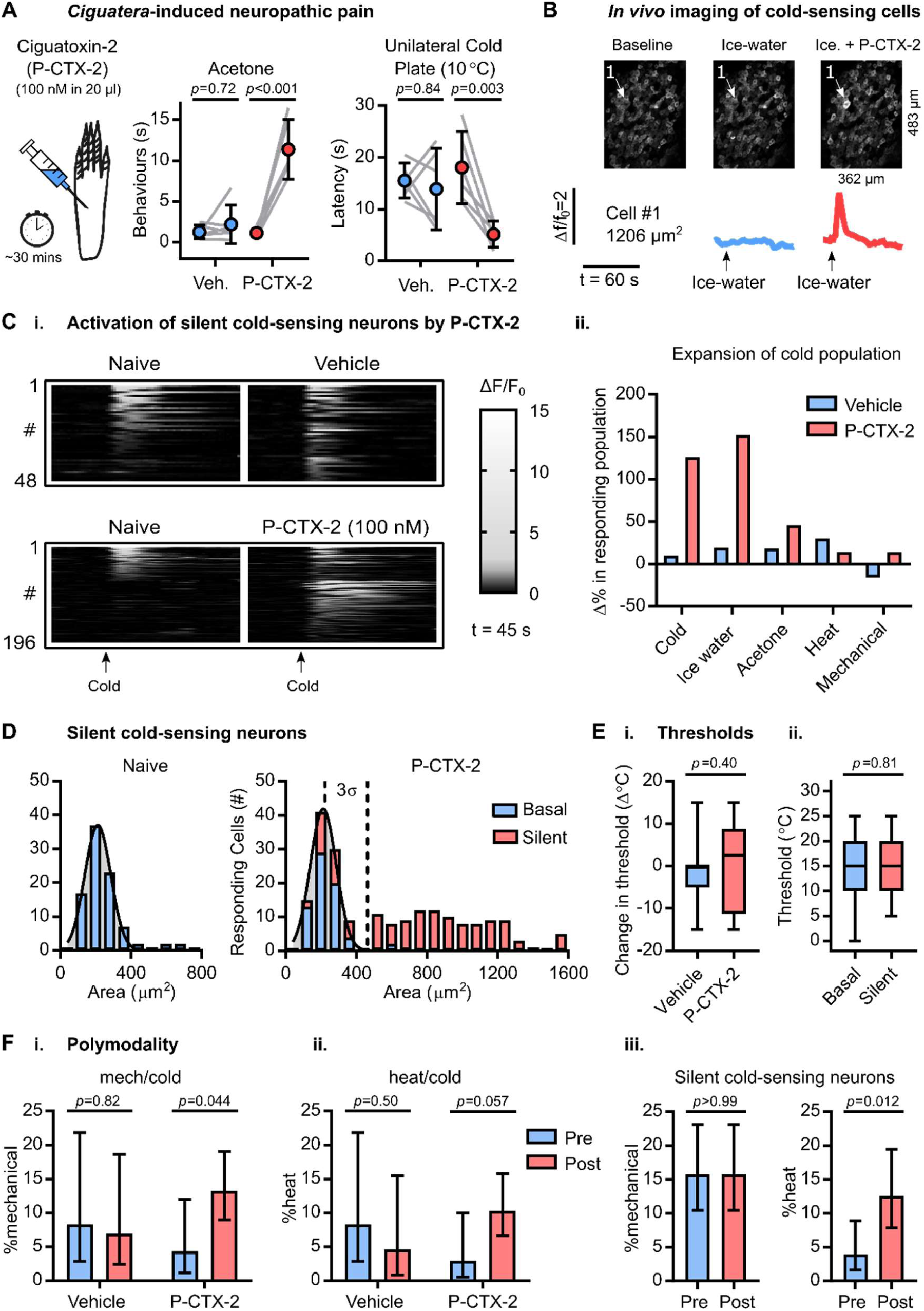
Silent cold-sensing neurons are activated by ciguatoxin-2. (A) Behavioural testing of the effect of 100 nM ciguatoxin-2 (P-CTX-2) on cold sensitivity n=6 for sham vehicle and n=6 for P-CTX-2. Means were compared by repeated measures 2-way ANOVA followed by post-hoc Sidak’s test. Error bars denote 95% confidence interval. (B) Example images and traces of a large-diameter neuron (Cell #1) that is basally cold-insensitive but begins to respond to cooling after treatment with P-CTX-2. (C) (i.) Heatmap showing the effect of P-CTX-2 on the number of neurons responding to a cold ice-water stimulus. n=48 for vehicle, and n=196 for P-CTX-2. (ii.) Summary of the change in the number of sensory neurons responding to each modality after treatment with P-CTX-2. (D) Histograms of cross-sectional area of all neurons responding to any cold stimulus in the naïve state (left, blue, n=91) and after P-CTX-2 (right). For P-CTX-2, blue denotes basally-responsive neurons that maintained their response to cold (n=70) and red denotes the silent cold-sensing neurons that were unmasked after treatment (n=136). The distribution of areas in the naïve state was fit by non-linear regression (least squares Gaussian; Bin width is 80 µm^2^; Mean=212.4 µm^2^, Std. Dev. 73.33 µm^2^). This model is plotted over the P-CTX-2 data to aid comparison with the dashed line denoting three standard deviations from the mean. The different in the distribution of areas between groups was assessed by Kolmogorov-Smirnov test (*p*<0.001). (E) (i.) Box plot of the change in activation threshold of basally cold-sensitive neurons before and after treatment with vehicle (n=35) or P-CTX (n=8). ii.) Box plot of the thermal activation threshold of all silent cold-sensing neurons unmasked by P-CTX-2 (n=43) compared to all cold-sensing neurons recorded from naïve mice (n=62). Medians were compared by Mann-Whitney test. (F) Quantification of the proportion of neurons responding ice-water that were also sensitive to either mechanical (i.) or heat (ii.) before and after treatment. Vehicle: n_pre_=36, n_post_=43. P-CTX-2: n_pre_=69, n_post_=174. (iii.) Comparison of the proportion of silent cold-sensing neurons that were responsive to other modalities before and after the induction of cold-sensitivity by P-CTX-2. n=127. The proportion of polymodal neurons was compared using χ^2^ test, and error bars denote 95% confidence intervals. For this experiment, 615 neurons responding to any stimulus either before or after treatment were recorded in 10 P-CTX-2-injected mice and 193 cells were recorded from 3 vehicle-injected animals.

We therefore imaged the change in sensory neuron cold sensitivity over the same time course (**Figure 3B**). After 30 minutes, P-CTX-2 transformed numerous cold-insensitive cells into neurons that showed robust responses to cooling (**Figure 3Ci**). As is evident in the heat-map, many basally-active cells actually lost their response to cold, however this was counterbalanced by the large number of cells that were unmasked by P-CTX-2, resulting in a net expansion of cold population, especially to ice-water (**Figure 3Cii**). The number of neurons responsive to any cold stimulus rose from 91 to 206 and for ice water went from 69 to 174. Compared to vehicle, there was no apparent effect of P-CTX-2 on the number of cells responding to other modalities (**Figure 3Cii**). The silent cold-sensing neurons unmasked by P-CTX-2 were generally large diameter with a mean cross-sectional area of 820.1 µm, in a manner clearly reminiscent of oxaliplatin and nerve injury (**Figure 3D**). The distribution of cell areas for heat was not markedly altered, although more small neurons responded to noxious pinch (**Figure S3A**).

How did P-CTX-2 treatment affect the properties of basally-active cold cells? We quantified the change in threshold of these cells after either P-CTX-2 or vehicle, and saw no difference (**Figure 3Ei**). Interestingly, P-CTX-2 reduced the peak cold response of these neurons compared to vehicle (**Figure S3B**). Silent cold-sensing neurons showed similar thermal activation thresholds to the basal population (**Figure 3Eii**), and their activity was not greater (**Figure S3C**). P-CTX-2 did not affect the peak response to other modalities (**Figure S3D**), however there was an increase in the fraction of polymodal mechano-heat neurons from 12% to 25% (**Figure S3E**).

P-CTX-2 increased mechano-cold polymodal neurons responding to ice-water from 4% to 13% (**Figure 3Fi**). Heat/cold polymodality was also enhanced, albeit not significantly (**Figure 3Fii**). Are polymodal silent cold-sensing neurons basally responsive to other modalities, or are both sensitivities acquired due to neuropathy? Interestingly, the proportion of identified silent cold-sensing neurons that responded to noxious mechanical stimuli was at 16% the same in the naïve state and after P-CTX-2. This indicates that at least some silent cold-sensing are responsive to noxious mechanical stimuli before the induction of neuropathy. On the other hand, few heat/cold cells showed a basal response to heat, indicating heat sensitivity is conferred by P-CTX-2 (**Figure 3Fiii**). These results were broadly similar when we looked only at the cold-sensing neurons defined by their response to acetone (**Figure S3F**).

These findings demonstrate cold allodynia induced by P-CTX-2 involves recruitment of silent cold-sensing neurons, some of which are basally sensitive to noxious pinch. Etiologically-distinct neuropathic pain states therefore give rise to cold pain by a similar mechanism of recruiting cold-insensitive sensory neurons to become cold-responsive. Given activation of silent cold-sensing neurons drives pain behaviours in mice and given their sensitivity to noxious mechanical stimulation, silent cold-sensing neurons can likely be classified as nociceptors.

### 2.4 Molecular characterization of silent cold-sensing neurons that drive cold allodynia in neuropathic pain

What is the molecular identity of silent cold-sensing neurons? To determine which sensory neuron subset they belong to, we crossed subset-specific Cre or CreERT2 mice with animals harbouring a Cre-dependent tdTomato reporter, on a Pirt-GCaMP3 background. This cross generated progeny expressing GCaMP3 in all sensory neurons but with tdTomato expression restricted to the cellular subset of interest (**Figure 4A**). Consequently, we were able to ask if functionally-identified silent cold-sensing neurons express molecular markers labelling major subpopulations of sensory neurons. We focused on oxaliplatin neuropathy because of its ease, reproducibility and clinical relevance. The percentage of neurons responding to any cold stimulus in vehicle and oxaliplatin-treated mice expressing each molecular marker is summarized in **Figure 4A**, with oxaliplatin split as before into basal and silent populations.

**Figure 4.**
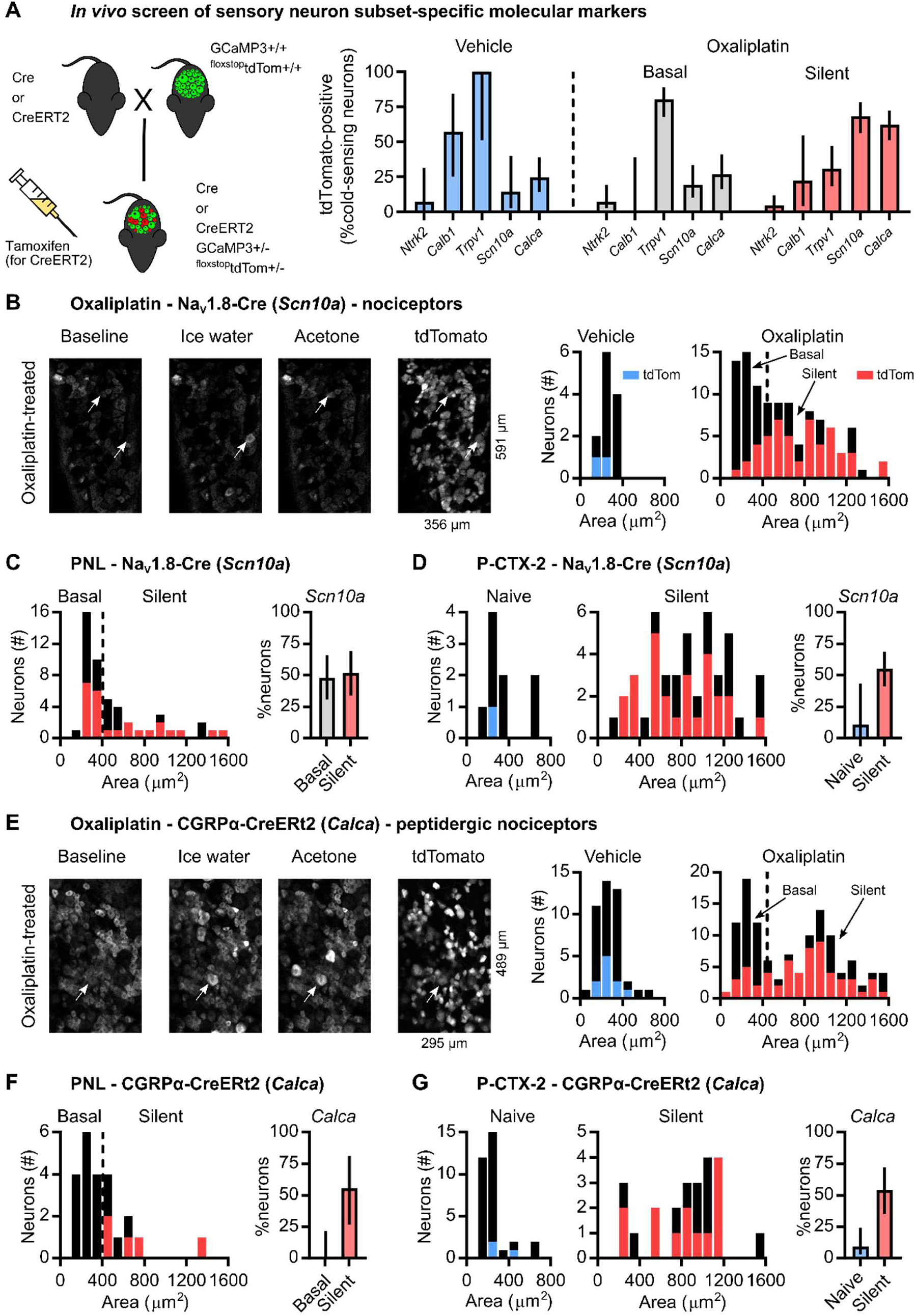
Silent cold-sensing neurons express peptidgergic nociceptor molecular markers Na_V_1.8 and CGRPα. (A) Cartoon (left) of breeding strategy used to generate GCaMP3 reporter mice for each subset of interest. Bar plot (right) showing overlap of reporter expression for each marker with functionally-defined cold-sensing neurons. TrkB-CreERT2 (*Ntrk2*): n_veh_=14 from 2 mice, n_oxa_=112 from 3 mice. Calb1-Cre (*Calb1*): n_veh_=7 from 1 mouse, n_oxa_=15 from 2 mice. Trpv1-Cre (*Trpv1*): n_veh_=4 from 1 mouse, n_oxa_=87 from 3 mice. Na_V_1.8-Cre (*Scn10a*): n_veh_=14 from 4 mice, n_oxa_=108 from 6 mice. CGRPα-CreERT2 (*Calca*): n_veh_=45 from 3 mice, n_oxa_=122 from 2 mice. (B) Example images (left) and histograms (right) showing overlap of Na_V_1.8-Cre-dependent tdTomato expression with cold-sensing neurons of different sizes in vehicle- and oxaliplatin-treated mice. Same data as in (A). (C) Histogram (left) and bar plot (right) showing overlap of Na_V_1.8-Cre-dependent tdTomato expression with different types of cold-sensing neurons in PNL-operated mice. n=50 cells from 1 mouse. (D) Histograms (left) and bar plot (right) showing overlap of Na_V_1.8-Cre-dependent tdTomato expression with basally-active and silent cold-sensing neurons in mice treated with P-CTX-2. n=56 cells from 4 mice. (E) Example images (left) and histograms (right) showing overlap of CGRPα-CreERT2-dependent tdTomato expression with cold-sensing neurons of different sizes in vehicle- and oxaliplatin-treated mice. Same data as in (A). (F) Histogram (left) and bar plot (right) showing overlap of CGRPα-CreERT2-dependent tdTomato expression with different types of cold-sensing neurons in PNL-operated mice. n=23 cells from 2 mice. (G) Histograms (left) and bar plot (right) showing overlap of CGRPα-CreERT2-dependent tdTomato expression with basally-active and silent cold-sensing neurons in mice treated with P-CTX-2. n=56 cells from 2 mice. Error bars denote 95% confidence intervals. As these data were obtained as part of an exploratory screen, no statistical hypothesis testing was performed.

Given their large size, we wondered if silent cold-sensing neurons might form part of low-threshold mechanoreceptor population. *Ntrk2* (TrkB-CreERT2) is a molecular marker for Aδ-fibre low-threshold mechanoreceptors and satellite glia cells, with glial and neuronal expression readily distinguished (**Figure S4A**). We found that just 4% of the silent cold-sensing neurons expressed *Ntrk2*, indicating they are unlikely to be Aδ-fibre low-threshold mechanoreceptors (**Figure 4A**). Silent cold-sensing neurons also showed minimal expression of the Aβ-fibre low-threshold mechanoreceptor marker *Calb*1 (Calb1-Cre) with only 22% overlap (**Figure 4A**).

In agreement with this, silent cold-sensing neurons that also responded to a stimulus that activates low-threshold mechanoreceptors were rarely observed, despite the possible mechanical confound associated with thermal stimulation (**Figure S5A**). When we imaged oxaliplatin-treated animals challenged with innocuous tactile stimuli, from 48 large-diameter silent cold-sensing neurons tested, not one responded to stroking the glabrous or hairy skin (with and against the grain) using a paint-brush or cotton swab (**Figure S5B**). In addition, when the glabrous skin was lightly stroked with a paint-brush, virtually no silent cold-sensing neurons unmasked by nerve injury (1/41) or ciguatera (2/136) responded to the low-threshold mechanical stimulus (**Figure S5B**). Both molecular marker expression and functional data therefore indicate silent cold-sensing neurons are unlikely to be low-threshold mechanoreceptors.

Interestingly, silent cold-sensing neurons showed 30% overlap with *Trpv1* lineage neurons (**Figure 4A**). The Trpv1-Cre line broadly labels a mixture of nociceptors and thermosensors, including basal cold-sensing neurons. This was recapitulated here, where essentially all small-diameter cold cells were marked by Trpv1-Cre, at 100% in vehicle, and 80% in the basal population following oxaliplatin.

Given that some silent cold cells respond to noxious mechanical stimuli, we hypothesized they express *Scn10a*, a marker of nociceptors.^33^ We previously showed that under physiological conditions the bulk of neurons sensing cold down to 0°C are negative for *Scn10a*, which encodes sodium channel Na_V_1.8.^20^ First, we surveyed the activity of *Scn10a*-positive neurons innervating the receptive field in response to different stimuli in naïve mice: the Na_V_1.8-Cre population encompassed heat- and mechanically-sensitive neurons of all sizes, but few cold-sensing cells (**Figure S4Bi**). In agreement with this, very few small-sized cold-sensing neurons were marked by Na_V_1.8-Cre in animals treated with either vehicle (14%) or oxaliplatin (19%). In contrast, we found 68% of the large-diameter silent cold-sensing neurons unmasked by oxaliplatin expressed Na_V_1.8 **(Figure 4A&B**). When we examined animals with cold allodynia evoked by PNL, 52% of silent cold-sensing neurons were marked by Na_V_1.8, although 48% of smaller cells also expressed Na_V_1.8 in this model (**Figure 4C**). Similarly, 55% of the silent cold-sensing neurons unmasked by P-CTX-2 were positive for Na_V_1.8, with only 11% overlap among the cold cells active in the naïve state (**Figure 4D**). *Scn10a* is widely expressed, labelling 55% of receptive field neurons in the naïve state, few of which, as noted, respond to cold (**Figure S4Bii**). However, the *Scn10a-*positive cold-sensing neurons unmasked during neuropathy are large in size and have cross-sectional areas markedly different to the typical distribution of areas observed for *Scn10a*-positive cells active in the naïve state (**Figure S4iii**). Thus, in all three neuropathic pain models tested here, a majority of silent cold-sensing neurons express Na_V_1.8, forming a unique subpopulation of large-diameter *Scn10a*-expressing presumptive nociceptors.

To determine which nociceptor subset silent cold-sensing neurons belonged to, we used the CGRPα-CreERT2 line to label *Calca*-expressing peptidergic nociceptors. In naïve mice, *Calca*-positive neurons were of small to medium size, mainly heat-sensitive and rarely responded to cold (**Figure S4Ci**). However, CGRPα-CreERT2 marked 62% of silent cold-sensing neurons in oxaliplatin neuropathy (**Figure 4A&E**). The silent cold-sensing neurons also overlapped substantially with *Calca*-expressing cells in mice with cold allodynia evoked by PNL (56%) or P-CTX-2 (54%), identifying silent cold cells as a set of peptidergic nociceptors commonly involved in cold allodynia (**Figure 4F&G**). Like *Scn10a*, the smaller basally-active cold cells were rarely positive for *Calca*. Although *Calca* is widely-expressed (**Figure S4Cii)**, the *Calca*-positive cold-sensing neurons unmasked by neuropathy had cross-sectional areas outwith the typical size distribution of responsive *Calca*-positive cells observed in naïve conditions (**Figure S4Ciii**). Thus, expression of *Scn10a* or *Calca* can be used to differentiate the silent and basal cold-sensing neurons. *Scn10a* and *Calca* are however expressed by a large and heterogenous set of nociceptors neurons; thus, these molecules are not selective markers for silent cold-sensing neurons.^34^

Do silent cold sensing neurons with a nociceptive identity contribute to behavioural cold hypersensitivity in neuropathic pain? We used diphtheria toxin to conditionally ablate silent cold-sensing neurons marked by Na_V_1.8-Cre to test their causal role in cold allodynia (**Figure 5A**). Note this ablation encompasses all cells expressing Na_V_1.8 and is not restricted to silent cold-sensing neurons. Imaging of mice where *Scn10a*-positive nociceptors are ablated shows very few of the large-diameter silent cold-sensing neurons are unmasked by oxaliplatin compared to Na_V_1.8-Cre mice lacking DTA (**Figure 5B**). The small basal cold-sensing neurons are retained after killing of *Scn10a*-positive neurons. Although nocifensive behaviour was not fully abolished, we observed a ∼50% decrease in oxaliplatin-evoked cold hypersensitivity in Na_V_1.8-Cre DTA animals (**Figure 5C**). The genetic identification and subsequent manipulation of silent cold-sensing neurons thus corroborates their causal contribution to cold allodynia in neuropathic pain.

**Figure 5.**
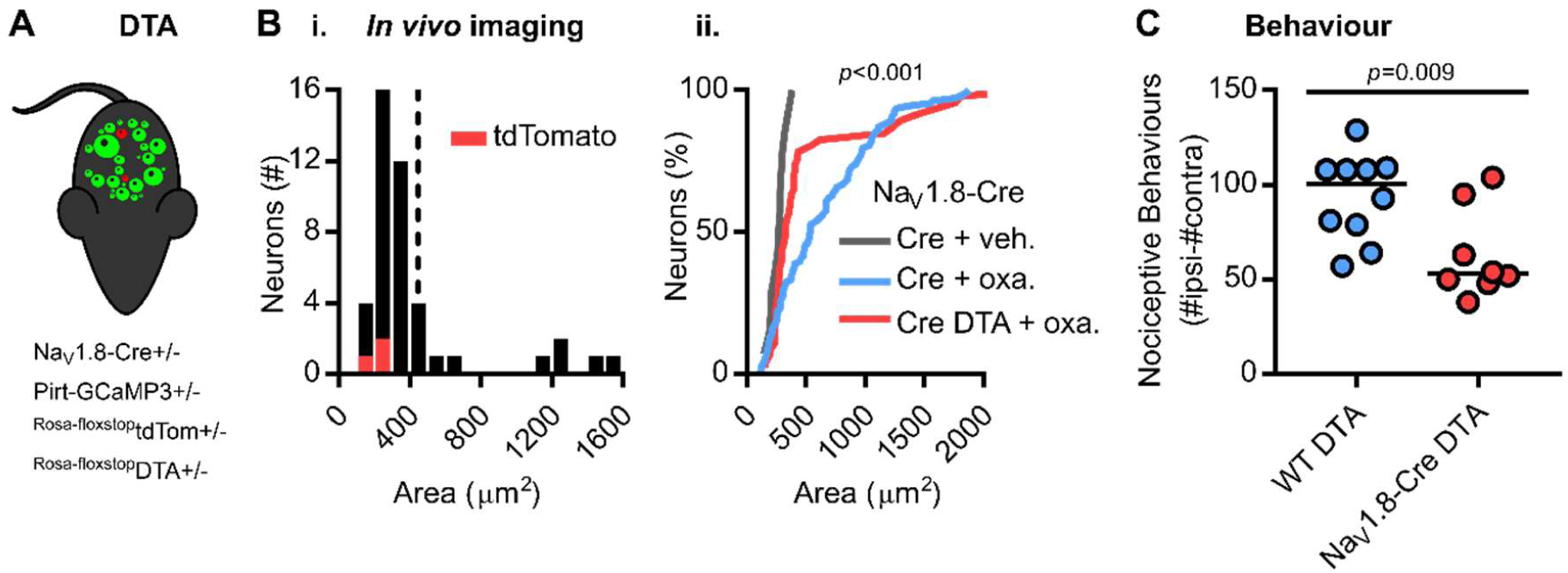
Diphtheria toxin-mediated ablation of Na_V_1.8-positive nociceptors decreases oxaliplatin-induced cold allodynia. (A) Cartoon of diphtheria toxin-mediated ablation of Na_V_1.8-positive neurons. (B) (i.) Histogram of cross-sectional areas of all cold-sensing neurons imaged in Na_V_1.8-Cre DTA mice treated with oxaliplatin. (ii.) Cumulative probability plot of cell areas in oxaliplatin-treated Na_V_1.8-Cre (blue) and Na_V_1.8-Cre DTA (red) mice, compared by Kolmogorov-Smirnov test. The distribution of cell areas in vehicle-treated Na_V_1.8-Cre mice is shown for comparison. n=108 cells from 6 oxaliplatin-treated Na_V_1.8-Cre mice, n=46 cells from 2 oxaliplatin-treated Na_V_1.8-Cre DTA mice, and n=14 cells from 4 vehicle-treated Na_V_1.8-Cre mice. (C) Quantification of the number of nociceptive behaviours in 5 minutes on the 5°C Cold Plate in 10 control and 8 Na_V_1.8-Cre DTA mice treated with oxaliplatin.

### 2.5 Molecular basis of cold detection by silent cold-sensing neurons

Enrichment of *Scn10a* and *Calca* in silent, but not basal, cold-sensing neurons offers a route to identifying molecules that might underlie pathological cold sensitivity. For mechanistic investigation, we focused on silent cold-sensing neurons responding to ice-water stimuli in the oxaliplatin model. Given that silent cold-sensing neurons express cold-resistant sodium channel isoform Na_V_1.8 and oxaliplatin is a known sodium channel activator, we wondered if Na_V_1.8 might be required for their response to cooling. Because Cre is knocked in directly at the Na_V_1.8 locus, homozygous Na_V_1.8-Cre mice lack both wild-type *Scn10a* alleles. TTX-resistant voltage-gated sodium currents could be recorded in medium to large-diameter Na_V_1.8-positive dorsal root ganglia neurons isolated from heterozygous, but not homozygous, Na_V_1.8-Cre reporter mice. This experiment demonstrates that homozygous Na_V_1.8-Cre mice are functionally Na_V_1.8 nulls, as well as indicating absent Na_V_1.9 expression in the cells recorded here (**Figure S6A**). We therefore treated homozygous Na_V_1.8-Cre mice expressing Cre-dependent tdTomato on a Pirt-GCaMP3 background with oxaliplatin. We observed numerous large-diameter silent cold-sensing neurons in null animals, many of which were positive for tdTomato (**Figure 6Ai**). There was no difference between oxaliplatin-treated mice heterozygous or homozygous for Na_V_1.8-Cre in the distribution of cross-sectional areas of cold-responsive cells (**Figure 6Aii**) or in the prevalence of tdTomato-expression in silent cold-sensing neurons (**Figure 6iii**). We also observed normal development of oxaliplatin-evoked cold allodynia in conventional Na_V_1.8 KO mice (**Figure S6B**). Imaging studies on Advillin-Cre conditional Na_V_1.7 KO mice expressing GCaMP3 also revealed intact recruitment of silent cold-sensing neurons by oxaliplatin (**Figure 6B**). Thus, pain-related sodium channels Na_V_1.8 and Na_V_1.7 are dispensable for silent cold-sensing neuron activity.

**Figure 6.**
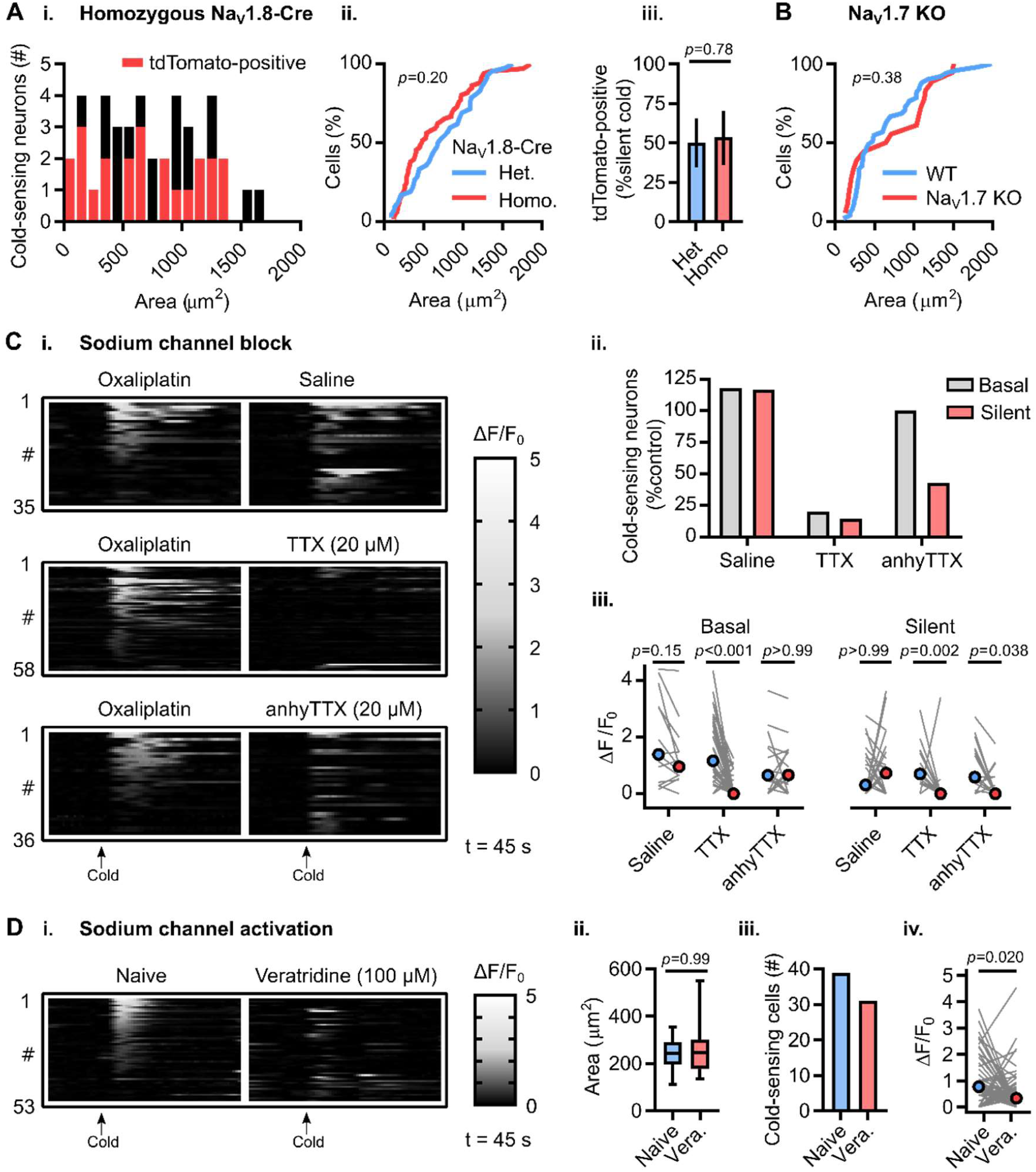
Voltage-gated sodium channel Na_**V**_1.6 is required for excitability, but is not sufficient for cold sensitivity, of silent cold-sensing neurons. (A) (i.) Histogram of cross-sectional areas of all cold-sensing neurons imaged in homozygous Na_V_1.8-Cre tdTomato mice with oxaliplatin. (ii.) Cumulative probability plot of cell areas in oxaliplatin-treated heterozygous and homozygous Na_V_1.8-Cre mice, compared by Kolmogorov-Smirnov test. (iii.) Bar plot showing the proportion of silent cold-sensing neurons expressing tdTomato in heterozygous and homozygous Na_V_1.8-Cre mice, compared using χ2 test. n=66 cells from 6 heterozygous and n=42 cells from 3 homozygous Na_V_1.8-Cre mice. (B) Cumulative probability plot of cell areas in oxaliplatin-treated WT and Na_V_1.7 KO mice, compared by Kolmogorov-Smirnov test. n=51 cells from 5 WT and n=18 cells from 2 Na_V_1.7 KO mice. (C) Heat maps (i.) and quantification (ii.) showing the effect of different sodium channel blockers on the number of basal and silent cold-sensing neurons in mice pre-treated with oxaliplatin. (iii.) Line plot showing the effect of blockers on median peak response, compared using Kruskall-Wallis test followed by Dunn’s multiple comparisons test. n=35 cells from 3 mice for saline, n=58 cells from 4 mice for TTX, n=36 cells from 2 mice for 4,9-anhydrous-TTX. (D) (i.) Heat map showing the effect of sodium channel activation with veratridine on the activity of cold-sensing neurons. (ii.) Box plot showing the size of cold-sensing cells is unaffected by veratridine, compared by Mann-Whitney test. (iii.) Bar plot showing veratridine reduces the number of cold-sensing neurons from 39 to 31. (iv.) Line plot showing veratridine reduces the response magnitude of cold-sensing cells, compared by Wilcoxon matched-pairs signed rank test. n=53 cells from 3 mice.

Which sodium channel isoform then is required for silent cold-sensing neuron excitability? Treatment of oxaliplatin-injected animals with TTX blocked activity in essentially all basal and silent cold-sensing neurons (**Figure 6C**). 4,9-anhydrousTTX, reported to preferentially inhibit Na_V_1.6, reduced the number of silent cold-sensing neurons by 57%. The effect of Na_V_1.6 blockade on basal cold-sensing neurons was comparable to saline, however (**Figure 6C**). Hence, Na_V_1.6 is likely the predominant sodium channel isoform in silent cold-sensing neurons. However, when we directly activated sodium channels with the pharmacological potentiator veratridine in naïve mice, we observed no unmasking of large-sized cells; indeed, the activity of cold-sensing neurons was paradoxically reduced (**Figure 6D**). Activation of sodium channels is therefore not sufficient to induce *de novo* cold sensitivity.

We have previously shown that basally cold-insensitive Na_V_1.8-positive neurons are enriched with *Kcna1* and *Kcna2*, which encode the voltage-gated potassium channels K_V_1.1 and K_V_1.2.^20^ These channels are thought to pass a voltage-dependent hyperpolarizing brake current that opposes depolarization evoked by cooling.^35^ Do these channels modulate the sensitivity of silent cold-sensing neurons to cooling? We hypothesized that pharmacological block of the I_KD_ current *in vivo* would unmask silent cold-sensing neurons. We imaged sensory neuron responses to cooling in Pirt-GCaMP3 mice at baseline and 30 minutes after intraplantar injection of the non-specific voltage-gated potassium channel blocker 4-aminopyridine (4-AP, 10 mM in 20 µl) (**Figure 7A**). 4-AP treatment triggered *de novo* sensitivity to cooling in previously cold-insensitive large diameter neurons (**Figure 7A-D**). The effect of 4-AP was reduced by pre-treatment with oxaliplatin (**Figure 7B-D**), suggesting that both 4-AP and oxaliplatin may act via the same cellular pathway.

**Figure 7.**
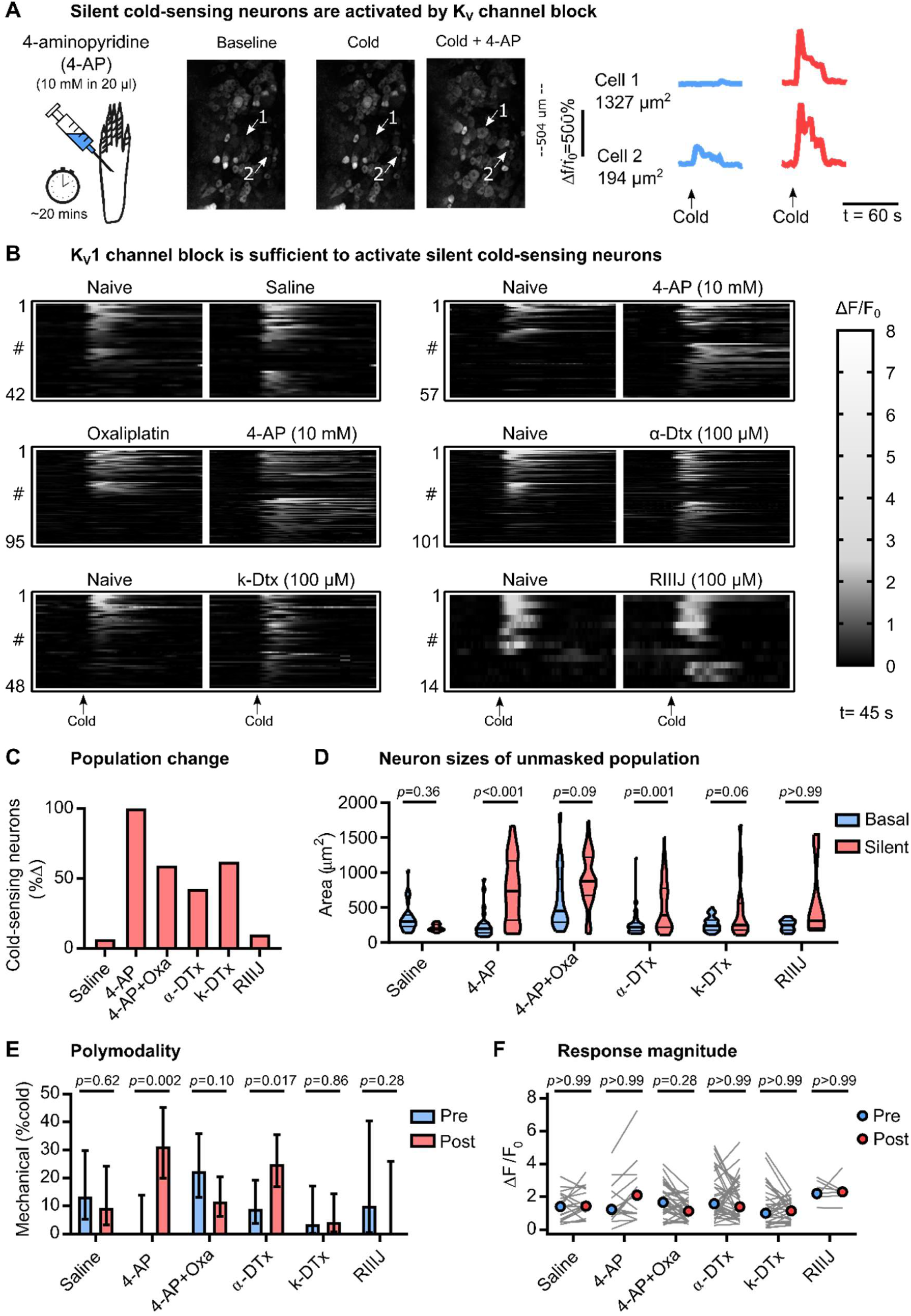
Blocking K_V_1.1 voltage-gated potassium channels is sufficient to induce *de novo* cold sensitivity in silent cold-sensing neurons. (A) Examples images and traces showing that peripheral blockade of voltage-gated potassium channels induces novel cold-sensitivity in normally cold-insensitive sensory neurons. (B) Heat maps showing the effect of different potassium channel blockers on the peripheral representation of cold. (C) Quantification showing the change in the number of cold-sensing neurons after treatment with different potassium channel blockers. (D) Violin plots showing the cross-sectional area of basal cold-sensing neurons in the naïve state (blue) and of silent cold-sensing neurons unmasked by potassium channel block (red). Medians were compared using Kruskall-Wallis test followed by Dunn’s multiple comparison test. (E) Bar plot of the percentage of polymodal cold-sensing neurons that also respond to noxious mechanical stimuli before (blue) and after (red) treatment with potassium channel blockers. Proportions were compared using a χ^2^ test. Error bars denote 95% confidence intervals. (F) No change in the median response magnitude of neurons that responded to cold both before (blue) and after (red) treatment with potassium channel blockers, as determined by Kruskall-Wallis test followed by Dunn’s multiple comparison test. n=42 from 3 saline-treated mice, n=57 from 6 4-AP-treated mice, n=95 from 3 4-AP-treated mice pre-injected with oxaliplatin, n=101 from 4 α-dendrotoxin-treated mice, n=48 from 3 k-dendrotoxin-treated mice and n=14 from 3 RIIIJ-treated mice.

Treatment with the selective K_V_1 antagonist α-dendrotoxin (100 µM) mimicked the effect of 4-AP (**Figure 7B-D**), indicating block of K_V_1 channels alone is sufficient to induce *de novo* cold sensitivity in silent cold-sensing neurons. k-dendrotoxin at 100 µM, a specific blocker of K_V_1.1, largely recapitulated the effect of α-dendrotoxin on silent cold-sensing neurons (**Figure 7B-D**). The K_V_1.2 blocker Conotoxin kM-RIIIJ (100 µM) had only minor effects (**Figure 7B-D**). 4-AP and α-dendrotoxin, but not k-dendrotoxin, increased the number of mechano-cold polymodal neurons (**Figure 7E**). Interestingly, none of the drug treatments significantly affected the activity of the basally-active population of cold neurons (**Figure 7F**). Overall, these data suggest that a functional reduction in K_V_1 channels, primarily mediated through K_V_1.1, can act a molecular switch that triggers *de novo* cold sensitivity in silent cold-sensing neurons and may therefore drive cold allodynia during neuropathic pain.

## 3 Discussion

The discovery of ion channels gated by cooling revolutionized our understanding of how sensory neurons detect cold in the healthy state.^7^ Thus far investigations into pathological cold pain have been hampered by our inability to monitor, in an objective fashion, the activity of cold-sensing neurons during chronic pain. *In vivo* imaging provides a unique opportunity to shed light on the cell and molecular changes causing cold allodynia in neuropathic pain. Here we show that neuropathic pain is associated with profound changes in the peripheral representation of cold, with large numbers of cold-insensitive peptidergic nociceptors becoming responsive to cooling and consequently driving pain.

### 3.1 Silent cold-sensing neurons in cold allodynia

Several mechanisms have been proposed to explain cold allodynia. Classical human experiments using A fibre blockade suggested that cold allodynia is caused by loss of Aδ cold fibres that tonically suppress C fibre nociceptor input.^36,37^ Cold allodynia could also result from central sensitization, with normal cold-evoked afferent activity driving pain through aberrant central circuits.^6^ Conversely, cold-sensitive nociceptors that usually respond to extreme cold may become activated to milder temperature drops and consequently trigger cooling-evoked pain.^27,32^ Finally, cold-insensitive nociceptors that provide noxious sensory input could acquire a *de novo* sensitivity to cooling.^10,23^

Using *in vivo* imaging, we found that activation of normally silent, large-diameter cold-sensing neurons is a common mechanism of cold allodynia in three clinically important, but etiologically-distinct, neuropathic pain states. In rat dorsal root ganglia, these large-diameter cells are neurofilament-rich Aβ and Aδ fibres that are mainly low-threshold mechanoreceptors, but also comprise many nociceptors.^38^ The silent cold-sensing neurons identified here typically had functional and molecular characteristics consistent with peptidergic A-fibre nociceptors that were observed in all three mouse models. Crucially we observed no changes in cold thresholds in the basally-active population of cold-sensing neurons. Thus, our findings suggest cold allodynia is a form of peripheral sensitization where a subpopulation of nociceptors gains an inappropriate sensitivity to the cold. This contrasts with tactile allodynia, which depends on peripheral drive from low-threshold mechanoreceptors expressing Piezo2.^39–41^

Electrophysiological studies support our conclusion that recruitment of cold-insensitive afferents is the primary driver of cold allodynia in the neuropathic pain models examined here. Recording of both rodent and human nerves exposed to oxaliplatin reveals myelinated A fibres fire more in the cold.^28,42,43^ Corroborating this, blockade of large fibres abolishes cold allodynia in humans with non-freezing cold injury and oxaliplatin neuropathy.^44,45^ Nerve injury likewise increases the fraction of cold-sensitive cutaneous sensory neurons.^27,46–48^ Treatment with ciguatoxin-1 also induced novel cold responses *in vitro*.^23^ There is a gradation, both in the severity of cold allodynia and in the number of silent cold-sensing neurons unmasked, evidenced by the much milder effects of our nerve injury model compared to oxaliplatin or ciguatoxin-2. Consistent with this, cold allodynia is more prevalent in patients with chemotherapy-induced neuropathy and ciguatera poisoning versus those with nerve injury.^3,4^ Nonetheless, because silent cold-sensing neurons are unmasked in all three neuropathic pain states tested here, these results do support a common underlying pathophysiology.

The dramatic expansion in cold-sensitivity after neuropathy contrasts with the sparse and modality-specific subpopulation of sensory neurons that signals cooling in the healthy state.^10–12^ We demonstrate that during neuropathy these ‘labelled lines’ conveying the sensation of cooling break down, with the emergence of large-diameter neurons that respond to both cold and noxious mechanical pinch, but rarely to heat or touch stimuli. Importantly, the percentage of silent cold-sensing neurons sensitive to pinch is likely an under-estimate due to the differing receptive field areas of these two stimuli. Modality-specific responses to sensory stimuli are widespread under normal conditions, but the number of polymodal neurons is increased by inflammatory mediators.^12^ Consequently, enhanced polymodality is a general feature of sensitized pain states.^49^

### 3.2 Molecular identity of silent cold-sensing neurons

Single cell RNA sequencing has created an increasingly detailed classification of sensory neurons subtypes.^50–52^ These studies confirm and extend traditional taxonomies based on electrophysiological, morphological and molecular markers.^38,53^ But to which of these subtypes do silent cold-sensing neurons belong?

Silent cold-sensing neurons unmasked by oxaliplatin did not express molecular markers for Aβ (*Calb1*) and Aδ (*Ntrk2*) LTMRs. This was surprising given that oxaliplatin preferentially modulates A fibre activity.^28,42,43,45^ However, even using a wide-range of light touch stimuli, applied to both hairy and glabrous skin, we were unable to evoke activity in silent cold-sensing neurons, so they are unlikely to be low-threshold mechanoreceptors. These findings are consistent with data which suggests TrkB-positive neurons are essential to tactile but not cold allodynia after nerve injury, indicating distinct modalities of allodynia are mechanistically different.^54,55^

The majority of silent cold-sensing neurons expressed Na_V_1.8, a classic marker of nociceptors, in all three tested models.^33^ This agrees with recent observations that Na_V_1.8, as well as labelling ∼90% of C fibre nociceptors, is also expressed by ∼40% of A fibre neurons, which also encompass nociceptors.^56^ Although Na_V_1.8 is not a selective marker of silent cold-sensing neurons, we found very few of the small diameter, basal cold-sensing cells express Na_V_1.8, consistent with previous observations.^20^ Importantly, DTA-mediated deletion of Na_V_1.8-positive neurons in normal mice had no effect on the moderate cold assays used here to examine allodynia. But after oxaliplatin treatment, deletion of Na_V_1.8-positive neurons including the newly unmasked cold sensors did result in the loss of cold allodynia, mechanistically linking the silent cold-sensing neurons with cold allodynia. Consistent with this, Na_V_1.8-DTA mice were previously shown to have deficient cold allodynia elicited by ciguatoxin-1, while deletion of HCN2 channels specifically in Na_V_1.8-positive neurons impairs cold allodynia in chronic constriction injury.^23,57^ Interestingly, Na_V_1.8 is also expressed by normally cold-insensitive cells that become responsive to cold after a 5 minute exposure to 1°C, hinting at a physiological role for the latent cold sensitivity of silent cold-sensing neurons.^20^

Because most silent cold-sensing neurons express CGRPα, they are likely to be peptidergic nociceptors, which comprise both A and C fibres. The bulk of CGRPα-positive sensory neurons also express Na_V_1.8.^34^ A recent *in vivo* imaging study of trigeminal ganglia showed that, following burn injury of the oral cavity, previously ‘silent’ neurons became newly sensitive to cooling.^10^ These neurons were also identified as peptidergic nociceptors on the basis of post-hoc immunohistochemical labelling for CGRPα.^10^ Tellingly, optogenetic inhibition of CGRPα-positive neurons transiently and reversibly relieves cold allodynia after spared nerve injury.^58^ Ciguatoxins also increase CGRP release and produce flare when injected into humans.^59,60^ Using *in vivo* imaging of the trigeminal ganglion, CGRPα-positive neurons were identified as a mixture of small-diameter polymodal nociceptors that respond to heat and large-diameter mechanonociceptors that respond to noxious mechanical stimulation of the hairy skin.^61^ This is in tune with our finding that large-diameter silent cold-sensing neurons often also respond to noxious pinch, although here our stimulation was restricted to the glabrous skin.

Taken together, these results identify silent cold-sensing neurons as mainly peptidergic nociceptors that express *Scn10a* and *Calca* molecular markers. Extending our previous observations that Na_V_1.8-positive sensory neurons signal prolonged and extreme cold, a further potential role for these nociceptors in mediating pathological responses to normal cooling is now apparent.^20^

### 3.3 Ionic mechanisms of *de novo* cold sensitivity

Our imaging data demonstrate that peripheral cold sensitivity is remarkably labile. By molecularly characterizing silent cold-sensing neurons, we aimed to identify the ion channels triggering the *de novo* cold responses in neuropathic pain. Both oxaliplatin and P-CTX-2 directly activate voltage-gated sodium channels.^28,30^ Sodium channels are also upregulated and redistributed after nerve injury.^62,63^ Which sodium channels are required for silent cold-sensing neuron excitability? Na_V_1.8 is expressed by silent cold-sensing neurons and is known to control action potential firing at low temperatures, but was dispensable for their excitability here.^16,20^ Given that oxaliplatin directly activates Na_V_1.6 and inhibition of Na_V_1.6 prevents oxaliplatin-evoked cold allodynia, Na_V_1.6 is the most probable subtype mediating cooling-evoked firing in silent cold-sensing neurons.^28,29^ We used intraplantar injection of 4,9-anhyTTX to test the role of Na_V_1.6, and saw a marked reduction in silent, but not basal, cold-sensing neuron activity, indicating Na_V_1.6 is the predominant sodium channel operating in silent cold-sensing neurons.

We have shown previously that voltage-gated potassium channels K_V_1.1 and K_V_1.2 were enriched in basally cold-insensitive, Na_V_1.8-positive neurons.^20^ Silent cold-sensing neurons therefore have high baseline expression of K_V_1 channels which pass a voltage-dependent excitability brake current opposing cold-induced depolarization.^18,27,64^ Blocking K_V_1 voltage-gated potassium channels with 4-aminopyridine or α-dendrotoxin consequently induced *de novo* cold sensitivity in silent cold-sensing neurons, and this effect was partially recapitulated by inhibiting K_V_1.1, but not K_V_1.2. Interestingly, activating sodium channels did not drive *de novo* cold-sensitivity, indicating that ectopic cold activation is not a consequence of a general increase in excitability, but specifically linked to eliminating K_V_1 channels. Indeed, K_V_1 channels are known to potently control action potential firing in response to sensory stimuli in both cold- and mechanically-sensitive nerve terminals.^18,65^

Does neuropathic pain lead to functional downregulation of K_V_1 channels? In dissociated cultures, 4-AP induces *de novo* cold sensitivity in neurons from control but not from mice with chronic constriction injury.^27^ This is paralleled by behavioural results which show 4-AP evoked cold hypersensitivity is suppressed in injured animals, indicating that nerve injury-induced cold allodynia operates via the same pathway as 4-AP to drive *de novo* cold sensitivity.^27^ Similarly, both oxaliplatin- and 4-AP-evoked cold allodynia can be blocked by inhibition of Na_V_1.6.^29^ Corroborating this, we found that 4-AP unmasked fewer silent cold-sensing neurons in mice pre-treated with oxaliplatin. The mechanism of K_V_1 channel of downregulation is unclear, and is likely to be specific to each disease state. Quantitative PCR of samples from oxaliplatin-treated mice reveal that there is a decrease in K_V_1.1 RNA, supporting transcriptional changes.^21^ Numerous reports have also found a decrease in both K_V_1.1 and K_V_1.2 expression following nerve injury.^66–72^ On the other hand, *in vitro* studies support a direct antagonist effect of both oxaliplatin and ciguatoxin on voltage-gated potassium channels.^43,73^ It is important to note that a causal link between peripheral neuropathy and K_V_1 channel activity was not investigated or explicitly demonstrated in our study.

K_V_1 channels are not intrinsically modulated by cooling so cannot provide the initial cold-induced depolarization in the peripheral nerve endings of silent cold-sensing neurons.^18,74^ Knockout and pharmacological data suggests the canonical cold transducers Trpm8 and Trpa1 are dispensable for the model of oxaliplatin neuropathy used here, although Trpa1 plays a role in cold allodynia evoked by ciguatoxin-1.^23,29^ This is consistent with our previous observation that Trpm8 is highly enriched in the Na_V_1.8-negative basally cold sensitive neurons, but is not expressed in the Na_V_1.8-positive silent cold-sensing neurons.^20^ Although the cold transducer in silent cold-sensing neurons is yet to be identified, the evidence indicates it is unlikely to be Trpm8. Closure of background potassium channels that maintain the hyperpolarized resting membrane potential is also proposed to mediate depolarization evoked by cooling.^75^ As several leak potassium channel subtypes have been implicated in cold allodynia, these channels are plausible candidate transducers.^21,26,76^

How do our findings inform the treatment of cold allodynia? An attractive therapeutic strategy would be to enhance or restore the hyperpolarizing voltage-gated potassium channel activity that normally prevents cold-evoked firing in the afferent terminals of silent cold-sensing neurons. The KCNQ voltage-gated potassium channel activators retigabine and flupirtine have been shown to reduce cold allodynia evoked by oxaliplatin and nerve injury.^77,78^ Selectivity of these compounds remains a problem and could be overcome using gene therapy. AAV-mediated over-expression of K_V_1.2 RNA impairs the development of cold allodynia induced by nerve injury in rats.^79^ The targeted over-expression of *Kcna1* in silent cold-sensing neurons would strongly ameliorate cold allodynia and thus represents a rational gene therapy for neuropathic pain.^80^

### 3.4 Conclusions

Overall, we show that cold allodynia results from a set of normally silent cold-sensing neurons gaining *de novo* cold sensitivity in neuropathic pain. Cold allodynia is therefore a form of peripheral sensitization. Silent cold-sensing neurons were identified as putative A-fibre peptidergic nociceptors based on their large diameter, response to noxious mechanical stimulation, and expression of molecular markers Na_V_1.8 and CGRPα. Silent cold-sensing neurons have basally high expression of K_V_1.1 and K_V_1.2 voltage-gated potassium channels that mediate an excitability brake current that opposes cooling-induced depolarization. Block of K_V_1 channels is sufficient to induce *de novo* cold sensitivity, pointing to downregulation of these channels during disease as a possible trigger of cold allodynia. By defining cells and molecules involved in cold allodynia, our findings will inform the development of better targeted therapeutics for neuropathic pain. The *in vivo* imaging data collected here provides a unique insight into the mechanisms underpinning cold allodynia, for the first time identifying silent cold-sensing neurons as critical drivers of cold-evoked neuropathic pain.

## Acknowledgements

We thank the Wellcome Trust (200183/Z/15/Z) and the MRC. DIM was supported by a PhD fellowship from the Wolfson Foundation. Versus Arthritis supported APL, QM and JNW with a programme grant (20200). We thank Xinzhong Dong for Pirt-GCaMP3 mice, David Ginty and members of the Molecular Nociception Group for Cre-lines, Richard Lewis for the gift of ciguatoxin-2, and Aida Marcotti for help with cold pain behaviour tests. We thank James Cox and other members of the Molecular Nociception Group for help and advice.

## Author contributions

DIM carried out the experiments with help from APL and QM and wrote the paper. ECE and JNW provided suggestions and reagents and contributed to the writing of the paper.

---

**Figure S1.**
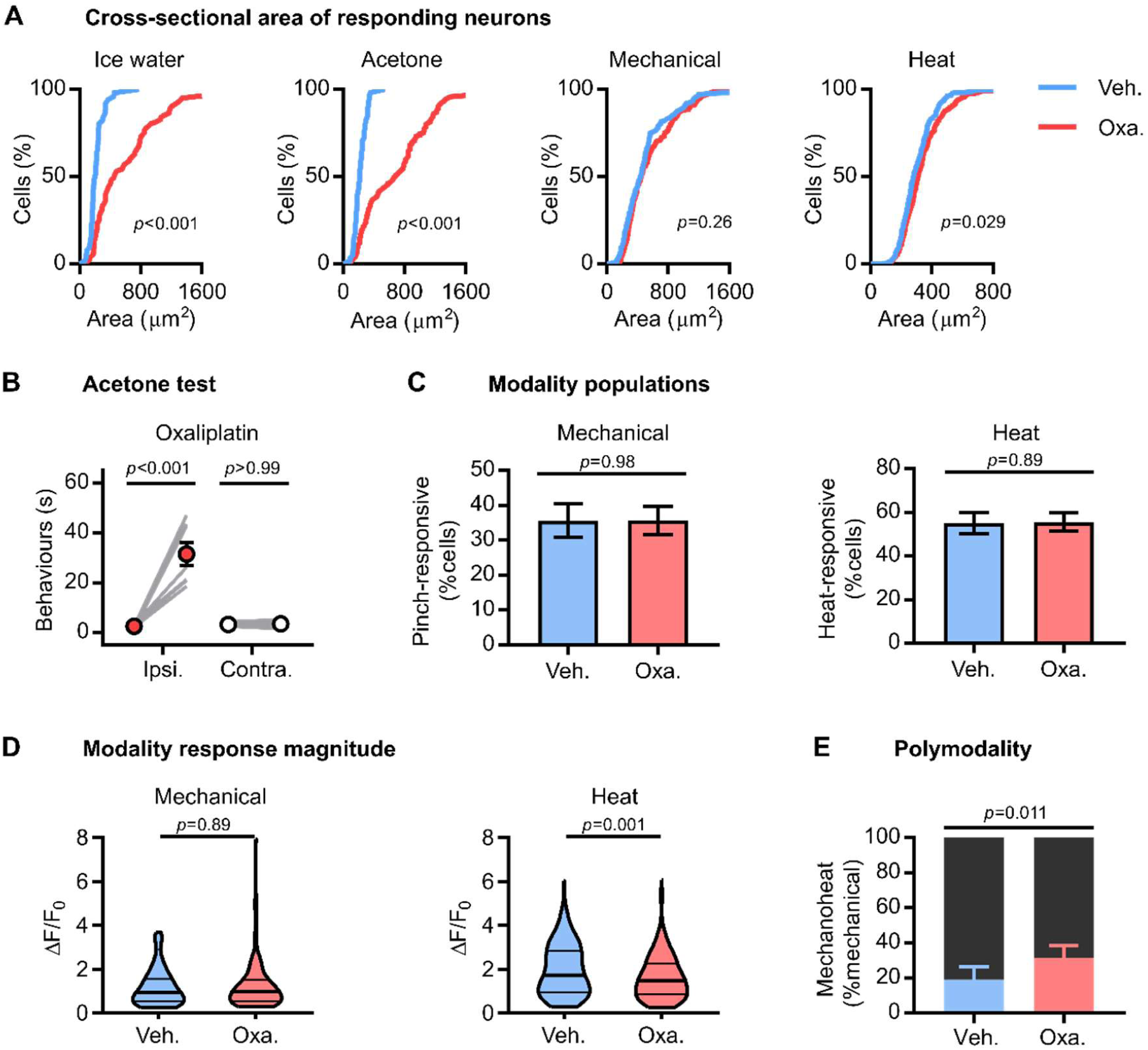
Modality-specific effects of oxaliplatin on sensory neuron physiology. (A) Cumulative probability plots of cross-sectional areas for cells responding to each stimulus modality, compared using Kolmogorov-Smirnov test. (B) Effect of oxaliplatin on acetone-evoked pain behaviour. Means before and after treatment were compared using repeated measures 2-way ANOVA with post-hoc Sidak’s test. n=7 for both. (C) Bar plots showing the proportion of all responding neurons responding to heat or mechanical stimuli, compared using χ^2^ test. Error bars denote 95% confidence interval. (D) Violin plots showing peak response evoked by each stimulus modality, compared using Mann-Whitney test. (E) Proportion of mechanically-sensitive neurons also responding to noxious heat, compared using χ^2^ test. Error bars denote 95% confidence interval. Ice-water: n_veh_=51, n_oxa_=81. Acetone: n_veh_=58, n_oxa_=145. Mechanical: n_veh_=136, n_oxa_=193. Heat: n_veh_=211, n_oxa_=301.

**Figure S2.**
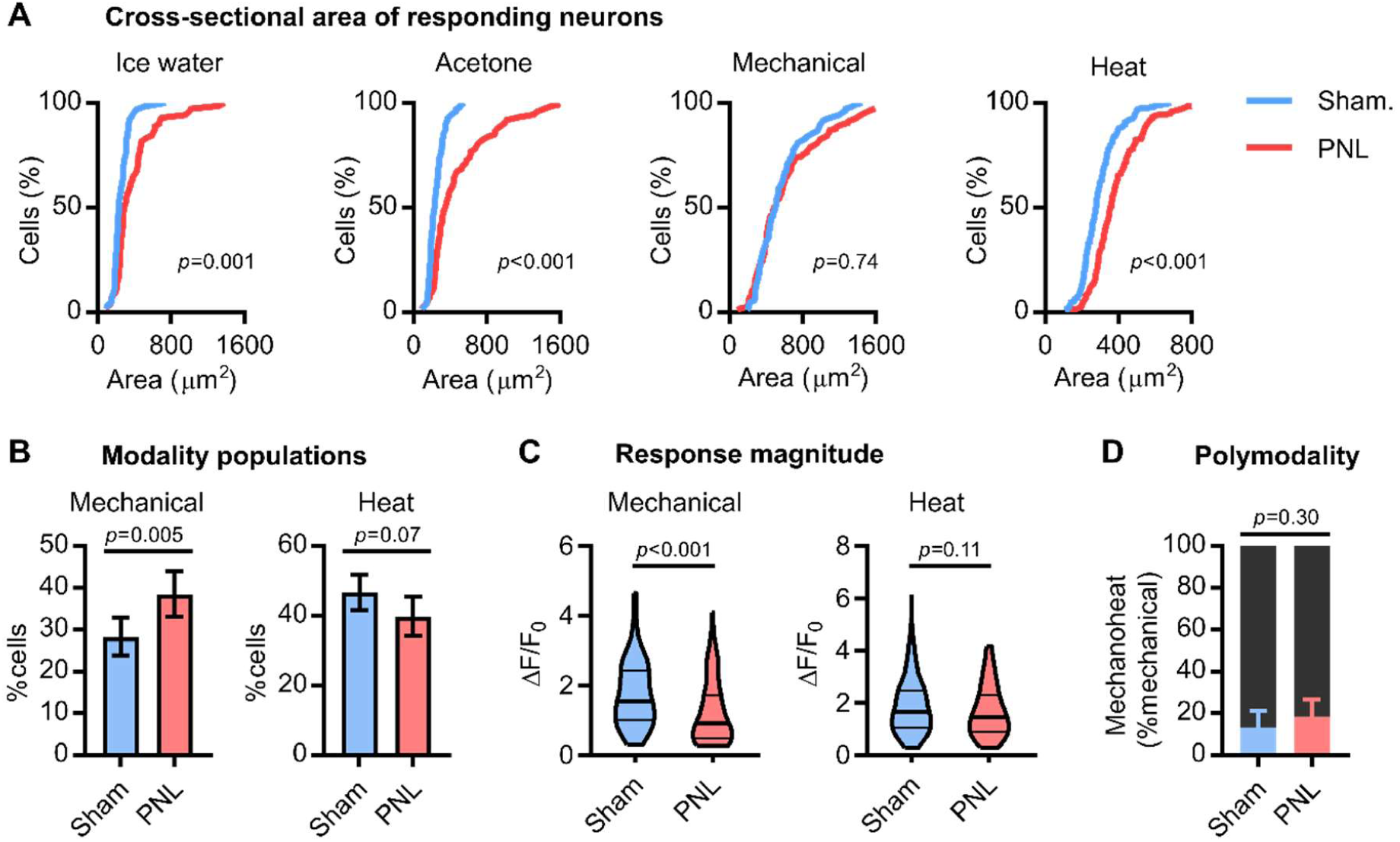
Modality-specific effects of partial sciatic nerve ligation on sensory neuron physiology. (A) Cumulative probability plots of cross-sectional areas for cells responding to each stimulus modality, compared using Kolmogorov-Smirnov test. (B) Bar plots showing the proportion of all responding neurons responding to heat or mechanical stimuli, compared using χ^2^ test. Error bars denote 95% confidence interval. (C) Violin plots showing peak response evoked by each stimulus modality, compared using Mann-Whitney test. (D) Proportion of mechanically-sensitive neurons also responding to noxious heat, compared using χ^2^ test. Ice-water: n_sham_=64, n_PNL_=71. Acetone: n_sham_=95, n_PNL_=73. Mechanical: n_sham_=105, n_PNL_=114. Heat: n_sham_=174, n_PNL_=118.

**Figure S3.**
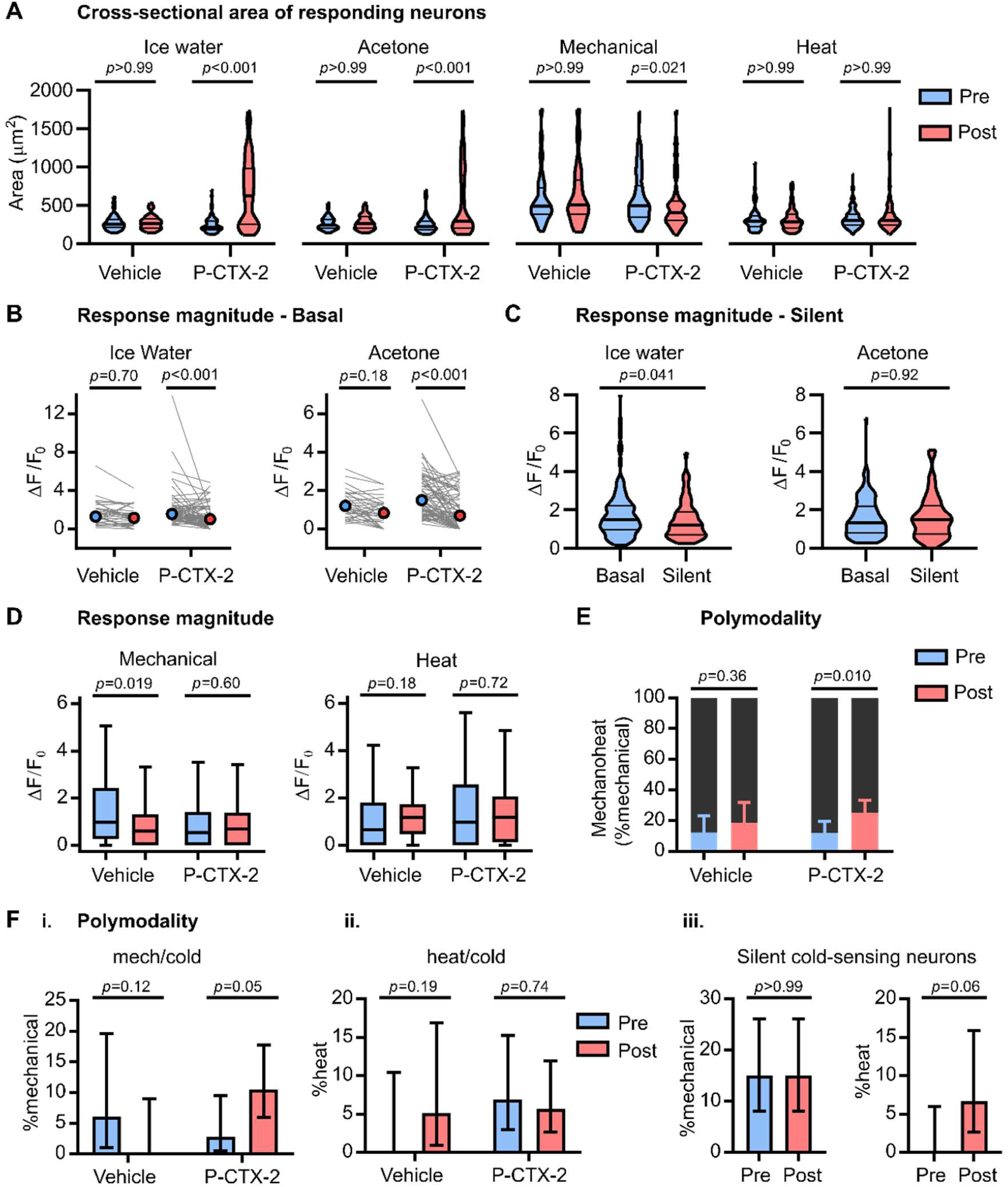
Modality-specific effects of ciguatoxin-2 on sensory neuron physiology. (A) Violin plots of cross-sectional areas for cells responding to each stimulus modality, compared using Kruskall-Wallis test followed by Dunn’s multiple comparisons test. (B) Line plots showing the median response magnitude of basally cold-sensitive neurons before and after treatment, compared using Kruskall-Wallis test followed by Dunn’s multiple comparisons test. (C) Violin plots showing the response magnitude of all silent cold-sensing neurons unmasked by P-CTX-2 (n=127 for ice-water, and n=60 for acetone) compared to all basally-active neurons recorded from naïve mice (n=105 for both). Medians were compared using Mann-Whitney test. (D) Box plots showing the median response magnitude of all mechanical and heat-responsive neurons before and after treatment. (E) Proportion of mechanically-sensitive neurons also responding to noxious heat, before and after treatment, compared using χ^2^ test. Error bars denote 95% confidence intervals. (F) Quantification of the proportion of neurons responding acetone that were also sensitive to either mechanical (i.) or heat (ii.) before and after treatment. (iii.) Comparison of the proportion of silent cold-sensing neurons activated by acetone that were responsive to other modalities before and after the induction of cold-sensitivity by P-CTX-2. n=60. The proportion of polymodal neurons was compared using χ^2^ test, and error bars denote 95% confidence intervals. Ice-water: vehicle: n_pre_=36, n_post_=43; P-CTX-2: n_pre_=69, n_post_=174. Acetone: vehicle: n_pre_=33, n_post_=39; P-CTX-2: n_pre_=72, n_post_=105. Mechanical: vehicle: n_pre_=57, n_post_=48; P-CTX-2: n_pre_=115, n_post_=131. Heat: vehicle: n_pre_=59, n_post_=77; P-CTX-2: n_pre_=211, n_post_=241.

**Figure S4.**
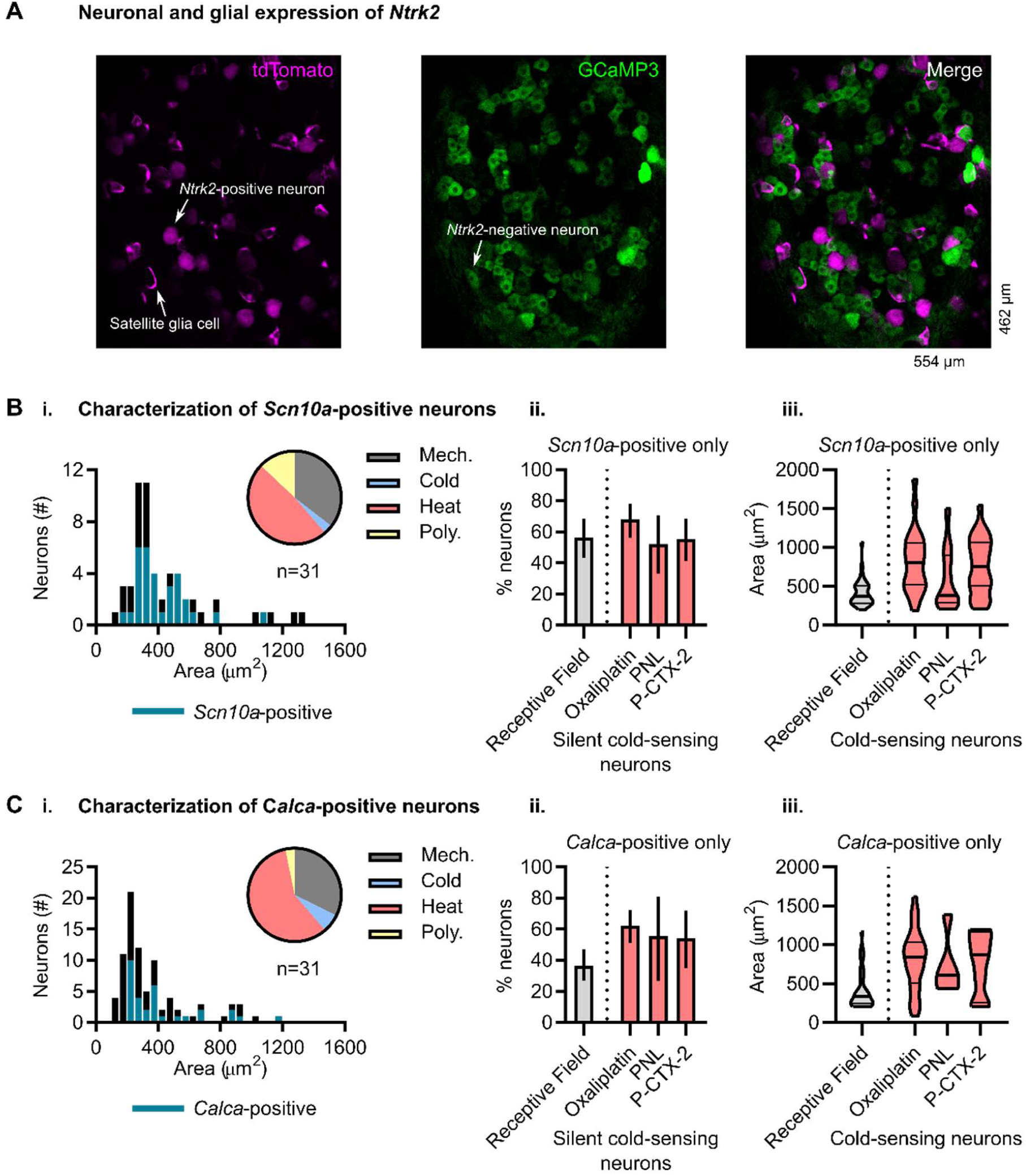
Functional and morphological characterization of sensory neurons expressing subset-specific molecular markers. (A) Representative *in vivo* images of the L4 dorsal root ganglion from a TrkB-CreERT2 mouse expressing Cre-dependent tdTomato and Pirt-driven GCaMP3. In the tdTomato channel (magenta), both neurons and satellite glia cells are observed labelled with tdTomato and thus likely express *Ntrk2*. In the GCaMP3 channel (green), only neuronal expression is seen. (B) (i.) In naïve mice, sensory neurons innervating the receptive field that also express Na_V_1.8-Cre (*Scn10a*) show a wide range of cross-sectional areas but are predominantly small to medium sized (histogram, magenta). These cells normally respond to noxious mechanical and heat stimuli but rarely to cold (inset, pie chart). Receptive field neurons are defined as all neurons responding to any stimulus applied to the plantar hindpaw. n=55 receptive field neurons from 4 mice, of which 31 are *Scn10a*-positive. (ii.) Comparison of the percentage of receptive field neurons that are *Scn10a*-positive in naïve mice compared to the percentage of silent cold-sensing neurons expressing *Scn10a* in different neuropathic pain models. n_RF_=55 cells from 4 mice. n_oxa_=66 silent cold-sensing neurons from 6 mice, n_PNL_=23 neurons from 1 mouse, n_P-CTX-2_=47 neurons from 4 mice. Errors denote 95% confidence intervals. (iii.) Comparison of the size of only *Scn10a*-positive receptive field neurons compared to all Scn10a-positive cold-sensing neurons in different neuropathic pain states. The *Scn10a*-positive cold-sensing neurons observed in neuropathy – the silent cold-sensing neurons – show cross-sectional areas outwith the typical distribution of areas for *Scn10a*-expressing receptive field neurons. n_RF_=31 cells from 4 mice. n_oxa_=53 *Scn10a*-positive cold-sensing neurons from 6 mice, n_PNL_=25 neurons from 1 mouse, n_P-CTX-2_=27 neurons from 4 mice. (C) (i.) In naïve mice, sensory neurons innervating the receptive field that express CGRPα-CreERT2 (*Calca*) show a wide range of cross-sectional areas but are predominantly small to medium diameter (histogram, magenta). These cells normally respond to noxious mechanical and heat stimuli but rarely to cold (inset, pie chart). Receptive field neurons are defined as all neurons responding to any stimulus applied to the plantar hindpaw. n=85 neurons from 2 mice. (ii.) Comparison of the percentage of receptive field neurons that are *Scn10a*-positive in naïve mice compared to the percentage of silent cold-sensing neurons expressing *Scn10a* in different neuropathic pain models. n_RF_=85 cells from 2 mice. n_oxa_=77 silent cold-sensing neurons from 2 mice, n_PNL_=9 neurons from 2 mice, n_P-CTX-2_=24 neurons from 2 mice. Errors denote 95% confidence intervals. (iii.) Comparison of the size of Scn10a-positive receptive field neurons compared to all Scn10a-positive cold-sensing neurons in different neuropathic pain states. The *Scn10a*-positive cold-sensing neurons observed in neuropathy – the silent cold-sensing neurons – show cross-sectional areas outwith the typical distribution of areas for *Scn10a*-expressing receptive field neurons. n_RF_=31 cells from 2 mice. n_oxa_=60 *Calca-*positive cold-sensing neurons from 2 mice, n_PNL_=5 neurons from 2 mice, n_P-CTX-2_=15 neurons from 2 mice.

**Figure S5.**
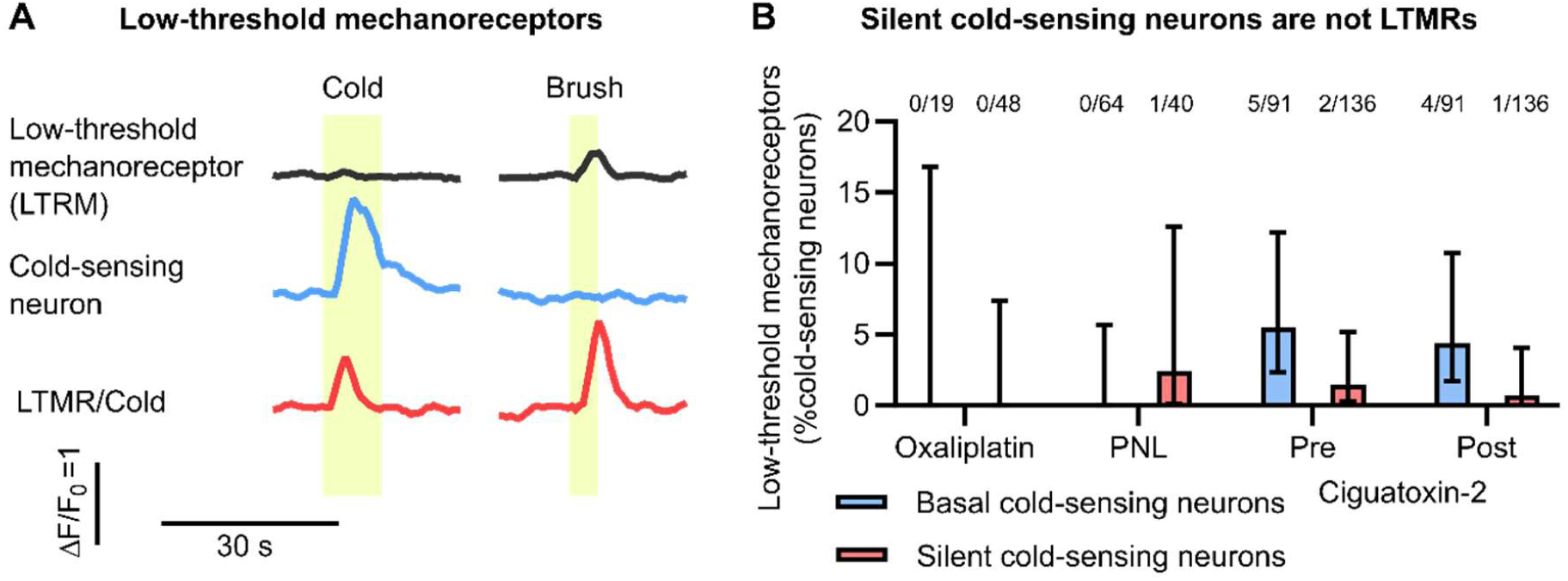
Silent cold-sensing neurons are not low-threshold mechanoreceptors. (A) Example traces of DRG neuron calcium signals in a naïve mouse in response to brush and cold stimuli. Example traces of a low-threshold mechanoreceptor (top) and a cold-sensing neurons (middle) are shown. A rare neuron responding to both brushing and cooling is also shown (bottom). (B) Bar plot showing the percentage of basal and silent cold-sensing neurons that show a response to low-threshold mechanical stimuli in each neuropathic pain model. For PNL and ciguatoxin-2, only brushing of glabrous skin was used; for oxaliplatin, cotton swap/brush was applied to glabrous skin and to hairy skin with and against grain. For ciguatoxin-2, the response to brush pre- and post-injection is shown. Across all models, only 3 out of 224 silent cold-sensing neurons showed a low-threshold mechanoreceptor type response.

**Figure S6.**
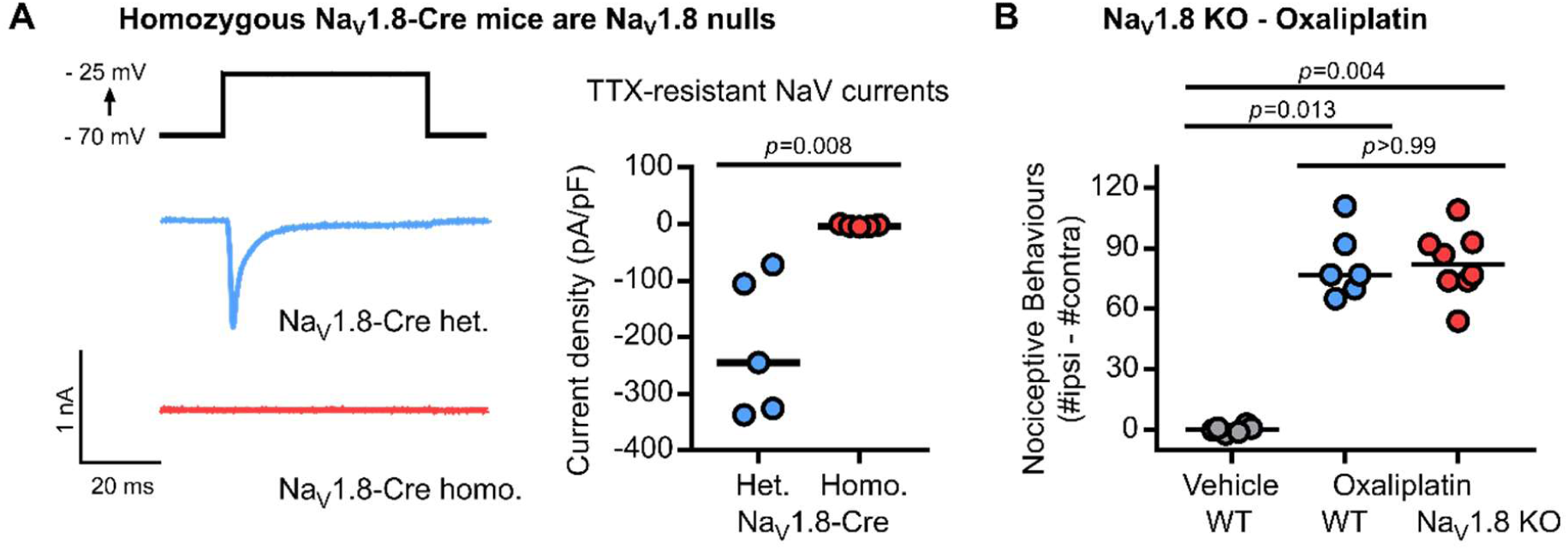
Mice lacking Na_**V**_1.8 develop oxaliplatin-induced cold allodynia. (A) Example traces and scatter plot of TTX-resistant sodium currents recorded from medium-sized DRG neurons cultured from heterozygous Na_V_1.8-Cre mice (blue). No TTX-resistant currents were observed in DRGs from homozygous Na_V_1.8-Cre mice (red). Median (line) current density was compared using the Mann-Whitney test. n=5 cells from 1 heterozygous mouse, and n=5 cells from 1 homozygous mouse. (B) Scatter plot of the effect of oxaliplatin on pain behaviours evoked by 5°C cold plate in WT and conventional Na_V_1.8 KO mice. Medians were compared using Kruskall Wallis test followed by Dunn’s multiple comparisons test. n=6 for WT treated with vehicle, n=6 for WT treated with oxaliplatin and n=8 for Na_V_1.8 KO treated with oxaliplatin.

## 4 STAR* Methods

### 4.1 Key Resources Table

#### 4.1.1 Experimental models: Organisms/Strains, Viruses, Pharmacological Agents

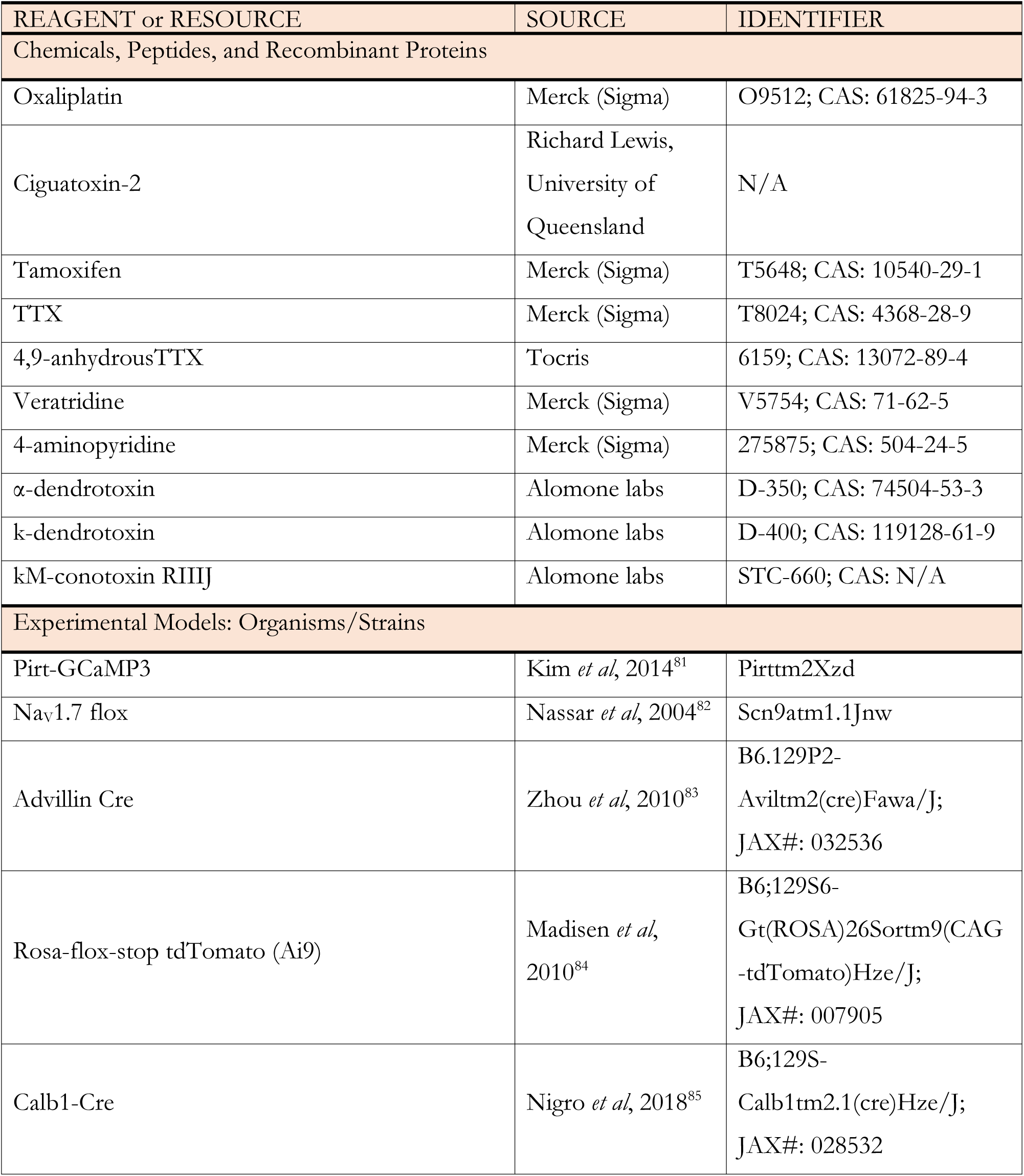
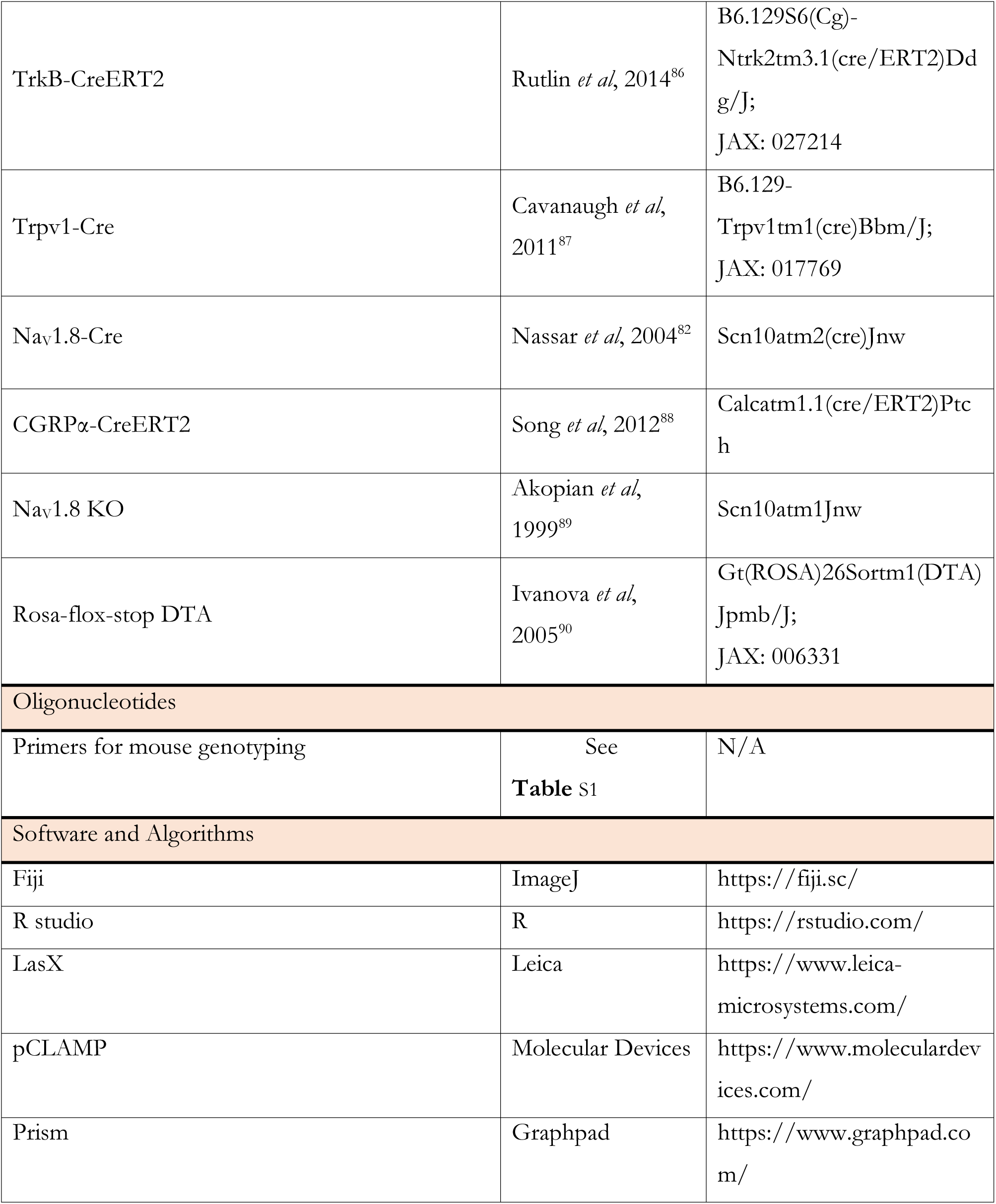

### 4.2 Lead contact and materials availability

Please direct any requests for resources and reagents to the Lead Contact, John N. Wood. This study did not generate any new unique reagents.

### 4.3 Experimental Model and Subject Details

#### 4.3.1 Animals

All animal procedures carried out at University College London were approved by University College London ethical review committees and conformed to UK Home Office regulations.

For experiments using wild-type mice, adult C57/BL6 mice from Charles River were used. Breeding strategies used to generate most of the mouse lines used in this study are as previously reported.^12,20,33,89,91^ Cre-dependent tdTomato reporter mice expressing GCaMP3 were generated by crossing subset-specific Cre mice with animals homozygous for Rosa-flox-stop tdTomato and homozygous for Pirt-GCaMP3. Na_V_1.8-Cre-dependent tdTomato and diphtheria toxin mice expressing GCaMP3 were generated by crossing Na_V_1.8 Cre mice with animals heterozygous for Rosa-flox-stop tdTomato, heterozygous Rosa-flox-stop DTA and homozygous for Pirt-GCaMP3. CGRPα-CreERT2 mice were given three 200 µl doses of a 1% tamoxifen solution on consecutive days between 6-8 weeks of age. TrkB-CreERT2 mice were given 200 µl doses of a 1% tamoxifen solution on consecutive days between 8-10 weeks of age. Tamoxifen was made-up in a 15% ethanol/85% sunflower oil.

Mice were housed on a 12:12 hour light-dark cycle with food and water available *ad libitum*. Adult (>6 weeks) mice were used for all studies. Both male and female animals were used for all experiments, in equal numbers where possible. These studies were not however designed to test for sex differences, and sexes were pooled for analysis. The number of animals used to generate each dataset is described in individual figure legends. For genotyping, genomic DNA was isolated from ear tissue or tail clip biopsy for PCR. Genotyping primers are summarized in Table S1.

**Table S1.**
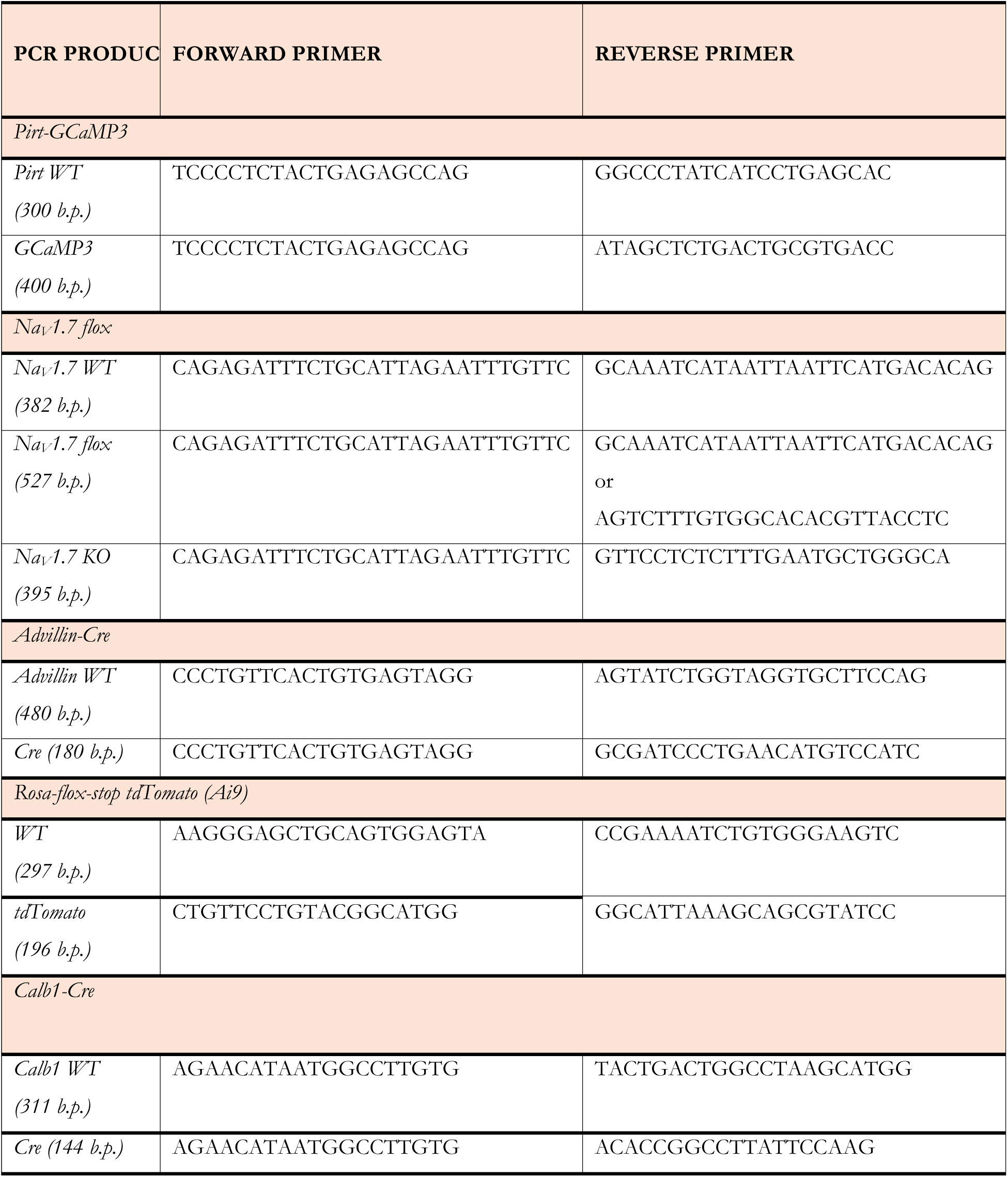
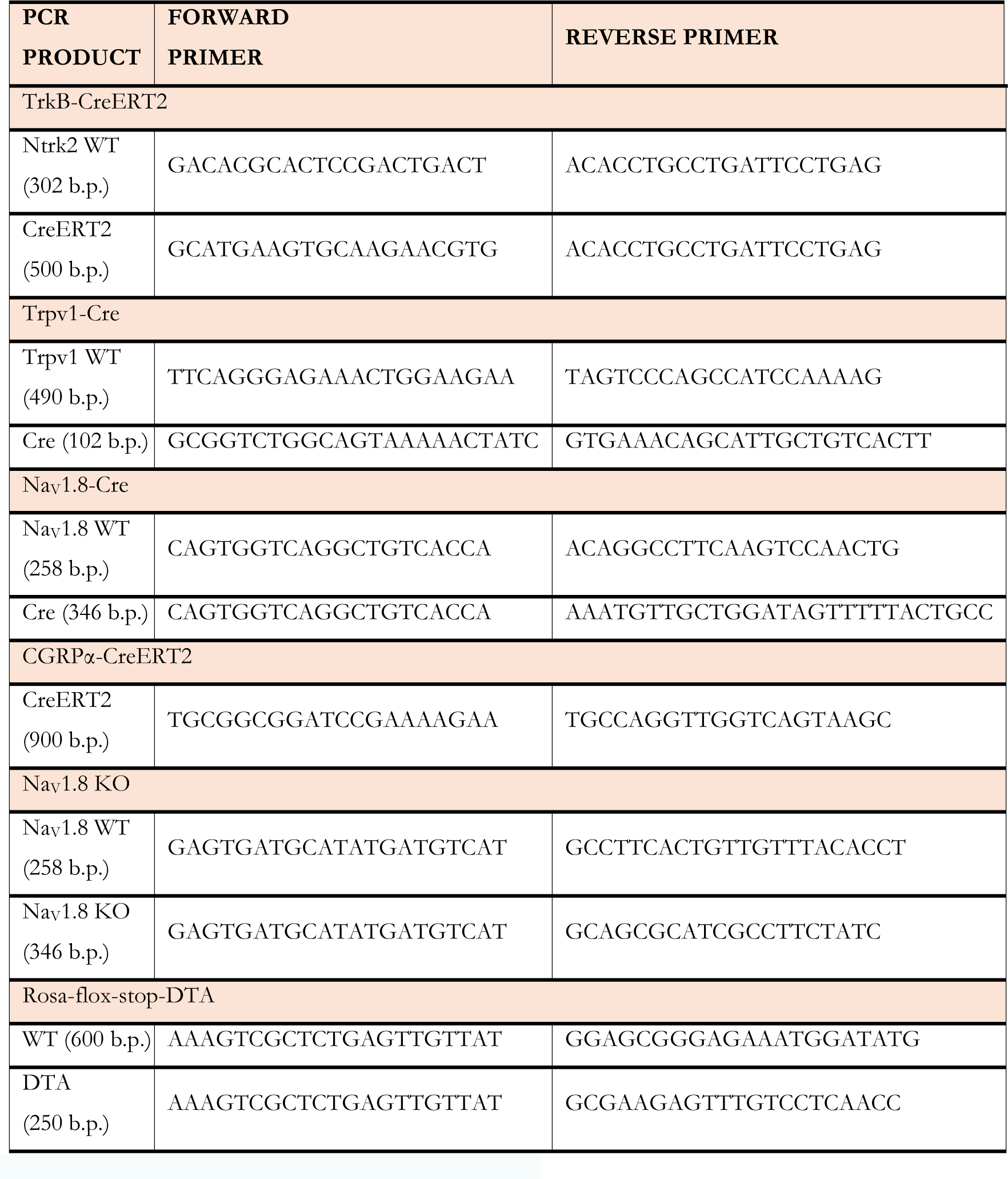
Summary of genotyping primers.

### 4.4 Method Details

#### 4.4.1 Neuropathic pain models

##### Oxaliplatin

Chemotherapy-induced neuropathy was studied in mice using the intraplantar oxaliplatin model first described by Deuis *et al* because this treatment recapitulates the rapid onset of cold allodynia in human patients infused with this drug.^29^ Oxaliplatin (Sigma) was made up in 5% glucose solution to an equivalent dose of 80 µg in 40 µl. This is because oxaliplatin is unstable in chloride-containing saline solution. Mice were treated by intraplantar injection into the left hindpaw. Vehicle was 5% glucose solution. Behavioural testing or imaging was assessed at least 3 hours after injection.

##### Partial Sciatic Nerve Injury

Peripheral nerve injury was studied in mice using a modified version of the Setlzer model.^92^ Surgical procedures were performed under isoflurane anaesthesia (2–3%) in adult mice. After the left thigh area was shaved and the skin sterilized with 70% ethanol, a longitudinal skin incision was made at the level of the femur. With the help of forceps, the muscle fibres were separated to allow visualization of the sciatic nerve. The partial nerve injury was induced by tying a tight ligature with 6-0 silk suture around approximately 1/3 of the diameter of the sciatic nerve. The skin was then closed with 6-0 Vicril suture and animals kept in a warm enclosure until complete recovery. Behavioural testing was performed at 2 and 4 weeks post-surgery, and imaging experiments were carried out between 4 and 5 weeks after surgery.

##### Ciguatoxin-2

Ciguatera poisoning was studied in mice using a modified version of the intraplantar ciguatoxin model first described by Vetter *et al* as this treatment isolates the sensory effects of the disease from the central and gastrointestinal symptoms.^23^ Ciguatoxin-2 (P-CTX-2) was a gift from Richard Lewis (University of Queensland). P-CTX-2 is highly-lipophilic and sticks to plastic surfaces. P-CTX-2 was therefore made up to 10 µM in 50% methanol solution, stored in a glass vial at −20°C and aliquoted using a metal/glass Hamilton syringe. The stock solution was diluted in saline containing 1% BSA to produce a final concentration of 100 nM. Mice undergoing imaging or behavioural testing were injected intraplantar with 20 µl of 100 nM P-CTX-2, and the effect of the drug measured 20-30 minutes after. Vehicle treatment was a 50% methanol solution diluted 100-fold in 1% BSA.

#### 4.4.2 *In Vivo* Calcium Imaging

##### Acquisition

Adult mice expressing GCaMP3 (6 to 14 weeks, male and female) were anesthetized using ketamine (100 mg/kg), xylazine (15 mg/kg) and acepromazine (2.5 mg/kg). Depth of anaesthesia was confirmed by pedal reflex and breathing rate. Animals were maintained at a constant body temperature of 37°C using a heated mat (VetTech). Lateral laminectomy was performed at spinal level L3-5. In brief, the skin was incised longitudinally, and the paravertebral muscles were cut to expose the vertebral column. Transverse and superior articular processes of the vertebra were removed using microdissection scissors and OmniDrill 35 (WPI). To obtain a clear image of the sensory neuron cell bodies in the ipsilateral dorsal root ganglion (DRG), the dura mater and the arachnoid membranes were carefully opened using microdissection forceps. The animal was mounted onto a custom-made clamp attached to the vertebral column (L1), rostral to the laminectomy. The trunk of the animal was slightly elevated to minimize interference caused by respiration. Artificial cerebrospinal fluid [containing 120 mM NaCl, 3 mM KCl, 1.1 mM CaCl_2_, 10 mM glucose, 0.6 mM NaH_2_PO_4_, 0.8 mM MgSO_4_, 1.8 mM NaHCO_3_ (pH 7.4) with NaOH] was perfused over the exposed DRG during the procedure to maintain tissue integrity, or the DRG was isolated by coating with silicone elastomer.

Images were acquired using a Leica SP8 confocal microscope. A 10x dry, 0.4-N.A. objective with 2.2 mm working distanced was used, with image magnification of 0.75-3x optical zoom. GCaMP3 was excited using a 488 nm laser line (1-15% laser power). tdTomato was excited using a 552 nm laser line (1-15% laser power). Filtering and collection of the emission light was optimized to maximize yield and minimize cross-talk (Leica Dye 164 Finer, LasX software, Leica). GCaMP was detected using a hybrid detector (100% gain) and tdTomato using a photomultiplier tube (500-600V gain). 512 x 512 pixel images were captured at a frame rate of 1.55 Hz, bidirectional scan speed of 800 Hz, and pixel dwell time of 2.44 µs.

Noxious and innocuous stimuli were applied to the left hindpaw, ipsilateral to the exposed DRG. For thermal stimuli, the paw was immersed with ice-water (nominally 0°C), acetone (100%) or water heated to 37°C or 55°C using a Pasteur pipette. For delivery of precise temperature stimuli, we used a Peltier-controlled thermode (Medoc). For mechanical stimuli, we used noxious pinch with serrated forceps, and innocuous brushing with a small paint-brush (ProArte-2) or cotton-swab. An interval of at least 30 s separated each stimulus application.

##### Pharmacology

TTX (20 µM in 20 µl saline for 20 minutes), 4,9-anhydro-TTX (20 µM in 20 µl saline for 20 minutes), 4-aminopyridine (10 mM in 20 µl saline for 20 minutes), α-dendrotoxin (100 µM in 20 µl saline for 20 minutes), k-dendrotoxin (100 µM in 20 µl saline for 20 minutes), conotoxin kappaM-RIIIJ (100 µM in 20 µl saline for 20 minutes) and veratridine (100 µM in 20 µl saline for 20 minutes) were all applied to the paw by intraplantar injection.

##### Analysis

Image stacks were registered to a reference image – typically, the first frame in the series – using the FIJI plugin TurboReg (accurate rigid body transformation) to correct for XY drift. Stacks that showed excessive Z movement were excluded from analysis. Regions of interest (ROI) were manually drawn around apparently responding cells using the free hand tool in FIJI. Mean pixel intensity over time for each ROI was extracted and analysed. The time series of mean pixel intensity for each ROI was smoothened by a four time point moving average to remove high-frequency noise. Next, we calculated the derivative of the mean pixel intensity. We calculated a mean baseline derivative for the 10 s preceding stimulus application. Neurons were classed as responders if, within 30 s of stimulus application, the maximum derivative was greater than the baseline derivative plus five standard deviations – that is, a Z-score of at least 5. We then calculated the ΔF/F_0_ value for each response to obtain a normalized measure of change in fluorescence. Neurons which showed a ΔF/F_0_ less than 0.25 were then discarded. Each trace was then manually screened as a further precaution against false positives. The remaining neurons that made up the responding population were then used for statistical analysis.

Cross-sectional area for each ROI in µm^2^ was also extracted in Fiji and analysed. Because nuclei are visible in the cells analysed here, we can justify comparing the areas of the cells present in the image as measures of their relative size.

The red channel of the reference image was used determine whether a cell was positive for tdTomato. Five regions of interest were drawn in background areas of the image clearly negative for tdTomato and average red fluorescence measured to calculate the mean and standard deviation of the background red fluorescence. Red fluorescence in responding cells was Z-scored versus the background value, and cells were counted as tdTomato positive if the Z-score was greater than 5.

#### 4.4.3 Behavioural Testing

All animal experiments were performed in accordance with Home Office Regulations. The investigator was blinded to treatment and/or genotype. Animals were acclimatized to handling and every effort was made to minimize stress during the testing. Both male and female animals were used.

##### Von Frey

Punctate mechanical sensitivity was measured using the up-down method of Chaplan to obtain a 50% withdrawal threshold.^93^ Mice were habituated for one hour in darkened enclosures with a wire mesh floor. A 0.4 g von Frey filament was applied to the plantar surface of the paw for 3 s. A positive response resulted in application of a filament of lesser strength on the following trial, and no response in application of a stronger filament. To calculate the 50% withdrawal threshold, five responses surrounding the 50% threshold were obtained after the first change in response. The pattern of responses was used to calculate the 50% threshold = (10[χ+Ϟδ])/10,000), where χ is the log of the final von Frey filament used, Ϟ = tabular value for the pattern of responses and δ the mean difference between filaments used in log units. The log of the 50% threshold was used to calculate summary and test statistics, in accordance with Weber’s Law.

##### Hot plate

The Hot Plate test measures supraspinal nociceptive behaviours in response to extreme heat.^94^ Mice were placed on the Hot Plate apparatus held at 50°C or 55°C. The tested ended when the animal showed a withdrawal behavior or licked its hindpaw. Cut-off time was 60 s for 50°C and 30 s for 55°C.

##### Cold Plate

Mice were placed on the Cold Plate apparatus.^95^ The Cold Plate was maintained at 5°C for 5 minutes while the animal was free to move around on the plate and the number of nociceptive behaviours (shaking, lifting, licking, guarding, biting) displayed by each paw were counted by the observer.^29^ In the unilateral Cold Plate test, animals were restrained, the ipsilateral paw was placed directly onto the plate maintained at 10°C and the time until paw withdrawal was then measured. This was repeated three times and a trial averaged response obtained.

##### Acetone Test

The acetone evaporation test measures the nociceptive behaviours triggered by evaporative cooling of the hindpaw.^96^ Mice were habituated for 1 hour in enclosures with a wire mesh floor. Using a home-made syringe, a 50 µl drop of acetone was applied to the ventral side of the ipsilateral hindpaw. The cumulative time where the ipsilateral hindpaw was engaged in nociceptive behaviours (lifting, shaking, licking, guarding, biting) over the ensuing 60 s was then counted. An average of three trials was obtained, with at least 10 minutes between trials.

#### 4.4.4 *In vitro* electrophysiology

Adult mice were killed by inhalation of a rising CO_2_ concentration followed by cervical dislocation to confirm death. Dorsal root ganglia (DRG) were dissected from the entire length of the spinal column and then digested in a pre-equilibrated enzyme mix for 45 minutes (37 °C, 5% CO_2_). The enzyme mix consisted of Hanks’ balanced salt solution containing collagenase (type XI; 5 mg/ml), dispase (10 mg/ml), HEPES (5 mM) and glucose (10 mM). DRGs were then gently centrifuged for 5 minutes at 300 revolutions per minute, the supernatant was discarded and replaced with warmed Dulbecco’s modified Eagle’s medium (DMEM), supplemented with L-glutamine (1%), glucose (4.5 g/litre), sodium pyruvate (110 mg/litre) and 10% fetal bovine serum (FBS). Next, DRGs were mechanically triturated with three fire-polished glass Pasteur pipettes of gradually decreasing inner diameter. Dissociated cells were then centrifuged again at 300 revolutions per minute, the supernatant was discarded and cells were re-suspended in the required volume of DMEM supplemented with FBS and nerve growth factor (50 ng/ml). Finally, cells were plated onto 12 mm glass coverslips coated with poly-L-lysine (1 mg/ml) and laminin (1 mg/ml). Cells were incubated at 37 °C in 5% CO_2_ and recordings were performed at room temperature 24 – 72 hours after dissociation.

Functional deletion of Na_V_1.8 was assessed by pharmacological isolation of TTX-resistant currents. Patch pipettes (tip resistance of 3-5 MΩ) were filled with intracellular solution containing: 140 mM CsF, 1 mM EGTA, 5 mM NaCl, 10 mM HEPES. Neurons were perfused with extracellular solution containing in: 70 mM NaCl, 70 mM Choline-Cl, 3 mM KCl, 1 mM MgCl_2_, 20 mM TEA-Cl, 0.1 mM CdCl_2_, and 10 mM Glucose. 5nM TTX was included in the extracellular solution to isolate TTX-resistant currents. Whole-cell recordings were obtained using an Axopatch 200B amplifier, filtered at 10 kHz and digitized at 50 kHz via a Digidata 1322A (Axon Instruments). tdTomato-expressing neurons from heterozygous and homozygous Na_V_1.8-Cre mice were voltage-clamped at −70 mV. Series resistance compensation was at least 60%. To measure the voltage-dependence of sodium channel activation, the holding command was dropped to −120 mV to de-inactivate all sodium channels and then a step-protocol from −80 to 20 mV was applied, in increments of 5 mV, to activate sodium channels.

### 4.5 Quantification and Statistical Analysis

For *in vivo* imaging experiments, n refers to the number of cells responding to any stimulus. For electrophysiology experiments, n refers to the number of recorded cells. For all imaging and physiology data, the number of animals used is indicated in the legend. For behavioural experiments, n refers to the number of animals. No power calculations were performed, however sample sizes are similar to those used in the field.

Datasets are presented using appropriate summary statistics as indicated in the legend, typically accompanied by raw data points or a representation of the underlying distribution. Behavioural data were generally assumed to be normal, as is typical in the field, and error bars denote mean ± 95% confidence interval. For *in vivo* imaging experiments, cells from all animals were pooled for analysis. Normality was not assumed when comparing cross-sectional areas or response magnitude of responding cells. These non-parameteric data are summarized using medians with quartiles, or cumulative probability plots. For categorical data, 95% confidence intervals around proportions were estimated using the Wilson-Brown method.

Tests of statistical comparison for each dataset are described in detail in figure legends. When comparing two groups, unpaired *t* test or Mann-Whitney test was used, depending on whether normality was assumed. When comparing the distribution of cell cross-sectional areas for two groups, the Kolmogorov-Smirnov test was used. For more than two groups, One-Way ANOVA or Kruskall-Wallis test was used with post-hoc tests corrected for multiple comparisons. When comparing the effect of two factors on multiple groups, a repeated-measures Two-Way ANOVA was used, with post-hoc tests corrected for multiple comparisons. For categorical data, proportions were compared using χ^2^ test. Curve fitting was performed using linear regression or non-linear regression functions.

Statistical tests were all performed using GraphPad Prism 7. An α-value of *p*=0.05 for significance testing was used. All p-values resulting from planned hypothesis testing are reported.

### 4.6 Data and code availability

Data are available from the lead contact on reasonable request.

## List of references

1. Breivik, H., Collett, B., Ventafridda, V., Cohen, R. & Gallacher, D. Survey of chronic pain in Europe: Prevalence, impact on daily life, and treatment. Eur. J. Pain 10, 287–287 (2006).

2. Jensen, T. S. & Finnerup, N. B. Allodynia and hyperalgesia in neuropathic pain: clinical manifestations and mechanisms. Lancet Neurol. 13, 924–935 (2014).

3. MacDonald, D. I., Wood, J. N. & Emery, E. C. Molecular mechanisms of cold pain. Neurobiology of Pain 7, 100044 (2020).

4. Yin, K., Zimmermann, K., Vetter, I. & Lewis, R. J. Therapeutic opportunities for targeting cold pain pathways. Biochem. Pharmacol. 93, 125–140 (2015).

5. Gold, M. S. & Gebhart, G. F. Nociceptor sensitization in pain pathogenesis. Nat. Med. 16, 1248–57 (2010).

6. Woolf, C. J. Evidence for a central component of post-injury pain hypersensitivity. Nature 306, 686–688 (1983).

7. Viana, F. & Voets, T. Heat Pain and Cold Pain. in The Oxford Handbook of the Neurobiology of Pain (Oxford University Press, 2019). doi: 10.1093/oxfordhb/9780190860509.013.13

8. Knowlton, W. M. et al. A sensory-labeled line for cold: TRPM8-expressing sensory neurons define the cellular basis for cold, cold pain, and cooling-mediated analgesia. J. Neurosci. 33, 2837–48 (2013).

9. Pogorzala, L. A., Mishra, S. K. & Hoon, M. A. The cellular code for mammalian thermosensation. J. Neurosci. 33, 5533–41 (2013).

10. Yarmolinsky, D. A. et al. Coding and Plasticity in the Mammalian Thermosensory System. Neuron 92, 1079–1092 (2016).

11. Wang, F. et al. Sensory Afferents Use Different Coding Strategies for Heat and Cold. Cell Rep. 23, 2001–2013 (2018).

12. Emery, E. C. et al. In vivo characterization of distinct modality-specific subsets of somatosensory neurons using GCaMP. Sci. Adv. 2, e1600990 (2016).

13. Bautista, D. M. et al. The menthol receptor TRPM8 is the principal detector of environmental cold. Nature 448, 204–208 (2007).

14. Colburn, R. W. et al. Attenuated cold sensitivity in TRPM8 null mice. Neuron 54, 379–86 (2007).

15. Dhaka, A. et al. TRPM8 is required for cold sensation in mice. Neuron 54, 371–8 (2007).

16. Zimmermann, K. et al. Sensory neuron sodium channel Nav1.8 is essential for pain at low temperatures. Nature 447, 855–8 (2007).

17. Lolignier, S. et al. The Nav1.9 Channel Is a Key Determinant of Cold Pain Sensation and Cold Allodynia. Cell Rep. 11, 1067–1078 (2015).

18. Madrid, R., de la Pena, E., Donovan-Rodriguez, T., Belmonte, C. & Viana, F. Variable Threshold of Trigeminal Cold-Thermosensitive Neurons Is Determined by a Balance between TRPM8 and Kv1 Potassium Channels. J. Neurosci. 29, 3120–3131 (2009).

19. Morenilla-Palao, C. et al. Ion Channel Profile of TRPM8 Cold Receptors Reveals a Role of TASK-3 Potassium Channels in Thermosensation. Cell Rep. 8, 1571–1582 (2014).

20. Luiz, A. P. et al. Cold sensing by NaV1.8-positive and NaV1.8-negative sensory neurons. Proc. Natl. Acad. Sci. U. S. A. 116, 3811–3816 (2019).

21. Descoeur, J. et al. Oxaliplatin-induced cold hypersensitivity is due to remodelling of ion channel expression in nociceptors. EMBO Mol. Med. 3, 266–78 (2011).

22. Nassini, R. et al. Oxaliplatin elicits mechanical and cold allodynia in rodents via TRPA1 receptor stimulation. PAIN® 152, 1621–1631 (2011).

23. Vetter, I. et al. Ciguatoxins activate specific cold pain pathways to elicit burning pain from cooling. EMBO J. 31, 3795–808 (2012).

24. Noël, J. et al. The mechano-activated K+ channels TRAAK and TREK-1 control both warm and cold perception. EMBO J. 28, 1308–18 (2009).

25. Alloui, A. et al. TREK-1, a K+ channel involved in polymodal pain perception. EMBO J. 25, 2368–76 (2006).

26. Pereira, V. et al. Role of the TREK2 potassium channel in cold and warm thermosensation and in pain perception. PAIN® 155, 2534–2544 (2014).

27. González, A. et al. Role of the Excitability Brake Potassium Current I KD in Cold Allodynia Induced by Chronic Peripheral Nerve Injury. J. Neurosci. 37, 3109–3126 (2017).

28. Sittl, R. et al. Anticancer drug oxaliplatin induces acute cooling-aggravated neuropathy via sodium channel subtype Na(V)1.6-resurgent and persistent current. Proc. Natl. Acad. Sci. U. S. A. 109, 6704–9 (2012).

29. Deuis, J. R. et al. An animal model of oxaliplatin-induced cold allodynia reveals a crucial role for Nav1.6 in peripheral pain pathways. Pain 154, 1749–57 (2013).

30. Inserra, M. C. et al. Multiple sodium channel isoforms mediate the pathological effects of Pacific ciguatoxin-1. Sci. Rep. 7, 42810 (2017).

31. Mckemy, D. D. Therapeutic potential of Trpm8 modulators. Open Drug Discov. J. 2, 81–88 (2010).

32. Chisholm, K. I., Khovanov, N., Lopes, D. M., La Russa, F. & McMahon, S. B. Large Scale In Vivo Recording of Sensory Neuron Activity with GCaMP6. eNeuro 5, (2018).

33. Abrahamsen, B. et al. The cell and molecular basis of mechanical, cold, and inflammatory pain. Science 321, 702–5 (2008).

34. Patil, M. J., Hovhannisyan, A. H. & Akopian, A. N. Characteristics of sensory neuronal groups in CGRP-cre-ER reporter mice: Comparison to Nav1.8-cre, TRPV1-cre and TRPV1-GFP mouse lines. PLoS One 13, e0198601 (2018).

35. Pertusa, M. & Madrid, R. The *I* KD current in cold detection and pathological cold pain. Temperature 4, 346–349 (2017).

36. Wahren, L. K., Torebjörk, E. & Jörum, E. Central suppression of cold-induced C fibre pain by myelinated fibre input. Pain 38, 313–319 (1989).

37. Yarnitsky, D. & Ochoa, J. L. Release of cold-induced burning pain by block of cold-specific afferent input. Brain 113, 893–902 (1990).

38. Lawson, S. N., Fang, X. & Djouhri, L. Nociceptor subtypes and their incidence in rat lumbar dorsal root ganglia (DRGs): focussing on C-polymodal nociceptors, Aβ-nociceptors, moderate pressure receptors and their receptive field depths. Current Opinion in Physiology 11, 125–146 (2019).

39. Eijkelkamp, N. et al. A role for Piezo2 in EPAC1-dependent mechanical allodynia. Nat. Commun. 4, (2013).

40. Murthy, S. E. et al. The mechanosensitive ion channel Piezo2 mediates sensitivity to mechanical pain in mice. Sci. Transl. Med. 10, eaat9897 (2018).

41. Szczot, M. et al. PIEZO2 mediates injury-induced tactile pain in mice and humans. Sci. Transl. Med. 10, eaat9892 (2018).

42. Kagiava, A., Tsingotjidou, A., Emmanouilides, C. & Theophilidis, G. The effects of oxaliplatin, an anticancer drug, on potassium channels of the peripheral myelinated nerve fibres of the adult rat. Neurotoxicology 29, 1100–1106 (2008).

43. Sittl, R., Carr, R. W., Fleckenstein, J. & Grafe, P. Enhancement of axonal potassium conductance reduces nerve hyperexcitability in an in vitro model of oxaliplatin-induced acute neuropathy. Neurotoxicology 31, 694–700 (2010).

44. Jørum, E. & Opstad, P.-K. A 4-year follow-up of non-freezing cold injury with cold allodynia and neuropathy in 26 naval soldiers. Scand. J. Pain 0, (2019).

45. Forstenpointner, J. et al. A-Fibers Mediate Cold Hyperalgesia in Patients with Oxaliplatin-Induced Neuropathy. Pain Pract. 18, 758–767 (2018).

46. Ji, G., Zhou, S., Kochukov, M. Y., Westlund, K. N. & Carlton, S. M. Plasticity in intact Ad- and C-fibers contributes to cold hypersensitivity in neuropathic rats. Neuroscience 150, 182–193 (2007).

47. Djouhri, L., Wrigley, D., Thut, P. D. & Gold, M. S. Spinal nerve injury increases the percentage of cold-responsive DRG neurons. Neuroreport 15, 457–60 (2004).

48. Xing, H., Chen, M., Ling, J., Tan, W. & Gu, J. G. TRPM8 Mechanism of Cold Allodynia after Chronic Nerve Injury. J. Neurosci. 27, 13680–13690 (2007).

49. Emery, E. C. & Wood, J. N. Somatosensation a la mode: plasticity and polymodality in sensory neurons. Current Opinion in Physiology 11, 29–34 (2019).

50. Usoskin, D. et al. Unbiased classification of sensory neuron types by large-scale single-cell RNA sequencing. Nat. Publ. Gr. 18, (2014).

51. Li, C.-L. et al. Somatosensory neuron types identified by high-coverage single-cell RNA-sequencing and functional heterogeneity. Cell Res. 26, 83–102 (2016).

52. Zeisel, A. et al. Molecular Architecture of the Mouse Nervous System. Cell 174, 999–1014.e22 (2018).

53. Emery, E. C. & Ernfors, P. Dorsal Root Ganglion Neuron Types and Their Functional Specialization. in The Oxford Handbook of the Neurobiology of Pain (Oxford University Press, 2018). doi: 10.1093/oxfordhb/9780190860509.013.4

54. Dhandapani, R. et al. Control of mechanical pain hypersensitivity in mice through ligand-targeted photoablation of TrkB-positive sensory neurons. Nat. Commun. 9, 1640 (2018).

55. Cobos, E. J. et al. Mechanistic Differences in Neuropathic Pain Modalities Revealed by Correlating Behavior with Global Expression Profiling. Cell Rep. 22, 1301–1312 (2018).

56. Shields, S. D. et al. Na v1.8 expression is not restricted to nociceptors in mouse peripheral nervous system. Pain 153, 2017–2030 (2012).

57. Emery, E. C., Young, G. T., Berrocoso, E. M., Chen, L. & McNaughton, P. A. HCN2 ion channels play a central role in inflammatory and neuropathic pain. Science 333, 1462–6 (2011).

58. Cowie, A. M., Moehring, F., O’Hara, C. & Stucky, C. L. Optogenetic Inhibition of CGRPα Sensory Neurons Reveals Their Distinct Roles in Neuropathic and Incisional Pain. J. Neurosci. 38, 5807–5825 (2018).

59. Touska, F. et al. Ciguatoxins Evoke Potent CGRP Release by Activation of Voltage-Gated Sodium Channel Subtypes NaV1.9, NaV1.7 and NaV1.1. Mar. Drugs 15, 269 (2017).

60. Zimmermann, K. et al. Analgesic treatment of ciguatoxin-induced cold allodynia. PAIN® 154, 1999–2006 (2013).

61. Ghitani, N. et al. Specialized Mechanosensory Nociceptors Mediating Rapid Responses to Hair Pull. Neuron 95, 944–954.e4 (2017).

62. Gold, M. S. et al. Redistribution of Na(V)1.8 in uninjured axons enables neuropathic pain. J. Neurosci. 23, 158–66 (2003).

63. Waxman, S. G., Kocsis, J. D. & Black, J. A. Type III sodium channel mRNA is expressed in embryonic but not adult spinal sensory neurons, and is reexpressed following axotomy. J. Neurophysiol. 72, 466–470 (1994).

64. Pertusa, M. & Madrid, R. The *I* KD current in cold detection and pathological cold pain. Temperature 4, 346–349 (2017).

65. Hao, J. et al. Kv1.1 Channels Act as Mechanical Brake in the Senses of Touch and Pain. Neuron 77, 899–914 (2013).

66. Ishikawa, K., Tanaka, M., Black, J. A. & Waxman, S. G. Changes in expression of voltage-gated potassium channels in dorsal root ganglion neurons following axotomy. Muscle and Nerve 22, 502–507 (1999).

67. Kim, D. S., Choi, J. O., Rim, H. D. & Cho, H. J. Downregulation of voltage-gated potassium channel alpha gene expression in dorsal root ganglia following chronic constriction injury of the rat sciatic nerve. Brain Res. Mol. Brain Res. 105, 146–52 (2002).

68. Park, S. Y. et al. Downregulation of voltage-gated potassium channel α gene expression by axotomy and neurotrophins in rat dorsal root ganglia. Mol. Cells 16, 256–259 (2003).

69. Rasband, M. N. et al. Distinct potassium channels on pain-sensing neurons. Proc. Natl. Acad. Sci. U. S. A. 98, 13373–13378 (2001).

70. Yang, E. K., Takimoto, K., Hayashi, Y., De Groat, W. C. & Yoshimura, N. Altered expression of potassium channel subunit mRNA and α-dendrotoxin sensitivity of potassium currents in rat dorsal root ganglion neurons after axotomy. Neuroscience 123, 867–874 (2004).

71. Calvo, M. et al. Altered potassium channel distribution and composition in myelinated axons suppresses hyperexcitability following injury. Elife 5, (2016).

72. Zhao, X. et al. A long noncoding RNA contributes to neuropathic pain by silencing Kcna2 in primary afferent neurons. Nat. Neurosci. 16, 1024–1031 (2013).

73. Birinyi-Strachan, L. C., Gunning, S. J., Lewis, R. J. & Nicholson, G. M. Block of voltage-gated potassium channels by Pacific ciguatoxin-1 contributes to increased neuronal excitability in rat sensory neurons. Toxicol. Appl. Pharmacol. 204, 175–186 (2005).

74. Viana, F., de la Peña, E. & Belmonte, C. Specificity of cold thermotransduction is determined by differential ionic channel expression. Nat. Neurosci. 5, 254–260 (2002).

75. Reid, G. & Flonta, M.-L. Cold transduction by inhibition of a background potassium conductance in rat primary sensory neurones. Neurosci. Lett. 297, 171–174 (2001).

76. Castellanos, A. et al. TRESK background K+ channel deletion selectively uncovers enhanced mechanical and cold sensitivity. J. Physiol. 598, 1017–1038 (2020).

77. Deuis, J. R. et al. Analgesic effects of clinically used compounds in novel mouse models of polyneuropathy induced by oxaliplatin and cisplatin. Neuro. Oncol. 16, 1324–32 (2014).

78. Abd-Elsayed, A. A. et al. KCNQ channels in nociceptive cold-sensing trigeminal ganglion neurons as therapeutic targets for treating orofacial cold hyperalgesia. Mol. Pain 11, 45 (2015).

79. Fan, L. et al. Impaired neuropathic pain and preserved acute pain in rats overexpressing voltage-gated potassium channel subunit Kv1.2 in primary afferent neurons. Mol. Pain 10, (2014).

80. Colasante, G. et al. CRISPRa-mediated Kcna1 upregulation decreases neuronal excitability and suppresses seizures in a rodent model of temporal lobe epilepsy. bioRxiv 431015 (2018). doi: 10.1101/431015

81. Kim, Y. S. et al. Central terminal sensitization of TRPV1 by descending serotonergic facilitation modulates chronic pain. Neuron 81, 873–87 (2014).

82. Nassar, M. A. et al. Nociceptor-specific gene deletion reveals a major role for Nav1.7 (PN1) in acute and inflammatory pain. Proc. Natl. Acad. Sci. U. S. A. 101, 12706–11 (2004).

83. Zhou, X. et al. Deletion of PIK3C3/Vps34 in sensory neurons causes rapid neurodegeneration by disrupting the endosomal but not the autophagic pathway. Proc. Natl. Acad. Sci. U. S. A. 107, 9424–9429 (2010).

84. Madisen, L. et al. A robust and high-throughput Cre reporting and characterization system for the whole mouse brain. Nat. Neurosci. 13, 133–140 (2010).

85. Nigro, M. J., Hashikawa-Yamasaki, Y. & Rudy, B. Diversity and connectivity of layer 5 somatostatin-expressing interneurons in the mouse barrel cortex. J. Neurosci. 38, 1622–1633 (2018).

86. Rutlin, M. et al. The cellular and molecular basis of direction selectivity of Aδ-LTMRs. Cell 159, 1640–51 (2014).

87. Cavanaugh, D. J. et al. Trpv1 Reporter Mice Reveal Highly Restricted Brain Distribution and Functional Expression in Arteriolar Smooth Muscle Cells. J. Neurosci. 31, 5067–5077 (2011).

88. Song, H. et al. Functional characterization of pulmonary neuroendocrine cells in lung development, injury, and tumorigenesis. Proc. Natl. Acad. Sci. U. S. A. 109, 17531–17536 (2012).

89. Akopian, A. N. et al. The tetrodotoxin-resistant sodium channel SNS has a specialized function in pain pathways. Nat. Neurosci. 2, 541–548 (1999).

90. Ivanova, A. et al. In vivo genetic ablation by Cre-mediated expression of diphtheria toxin fragment A. Genesis 43, 129–135 (2005).

91. Minett, M. S. et al. Distinct Nav1.7-dependent pain sensations require different sets of sensory and sympathetic neurons. Nat. Commun. 3, 791 (2012).

92. Seltzer, Z., Dubner, R. & Shir, Y. A novel behavioral model of neuropathic pain disorders produced in rats by partial sciatic nerve injury. Pain 43, 205–218 (1990).

93. Chaplan, S. R., Bach, F. W., Pogrel, J. W., Chung, J. M. & Yaksh, T. L. Quantitative assessment of tactile allodynia in the rat paw. J. Neurosci. Methods 53, 55–63 (1994).

94. Woolfe, G. & MacDonald, A. D. The evaluation of the analgesic action of pethidine hydrochloride (Demerol). J. Pharmacol. Exp. Ther. 80, (1944).

95. Allchorne, A. J., Broom, D. C. & Woolf, C. J. Detection of cold pain, cold allodynia and cold hyperalgesia in freely behaving rats. Mol. Pain 1, (2005).

96. Yoon, C., Young Wook, Y., Heung Sik, N., Sun Ho, K. & Jin Mo, C. Behavioral signs of ongoing pain and cold allodynia in a rat model of neuropathic pain. Pain 59, 369–376 (1994).

